# Light stimulation into dorsal raphe nucleus contributes to antidepressant effect for a stressed rat model

**DOI:** 10.1101/821421

**Authors:** Xiaotao Li

## Abstract

**Background:** Light therapy is frequently demonstrated by clinical trials to be effective to seasonal or non-seasonal major depression. However, the pathway underlying the light effect on mood remains unclear. Since a retino-raphe pathway was previously indicated to modulate 5-HT production, we hypothesize that the retinal projection into dorsal raphe nucleus (DRN) may play an important role in the light therapy for depression.

**Methods:** A rat model of 14-day corticosterone administration (40 mg/kg/day subcutaneous injection) was mainly used to test the effect of light therapy on non-seasonal depressant-like behavior, and the involved neural circuitry and neurochemistry as well.

**Results:** Behavior results revealed that the bright light therapy especially with the blue light of 470 nm and 400 lux, effectively reversed the depression-like responses in those stressed rats. After elimination of retino-raphe projection using immunotoxin (Saporin) the effect of light therapy was significantly attenuated. Whereas activation of retino-raphe projection using HM3q chemogenetics was shown an effect similar to fluoxetine treatment. Furthermore, 5-HT3A positive GABA cells in the DRN were activated with high c-Fos expression that involved in an inhibition of 5-HT synthesis and a subsequent depressive behavior. While light therapy through retino-raphe projection deactivated the hyperaction of those GABA cells in the DRN; that eventually contributed to the antidepressant effect from light therapy.

**Conclusions:** Our results indicate that the retino-raphe circuitry engaged antidepressant effect in DRN that contributed to the light therapy to the non-seasonal depression. 5-HT3A positive GABA cells in DRN was indicated to mediate this function of retino-raphe projection.

## Introduction

Light is sufficiently powerful to exert an effect on affective states [1, 2]. Such as the duration of sunshine significantly modulated the mood of healthy people through serotonergic activity in their brains [3, 4]. Blue-enriched artificial light in the workplace was able to improve subjective alertness and performance [5, 6]. Moreover, recently clinical trials demonstrated that light therapy was not only effective for seasonal affective disorder (SAD, namely seasonal depression) [7], but also useful for non-seasonal depression (in which major depressive episode is not obviously associated with seasonal pattern) [8, 9]. Light therapy shows its promising in the treatment of depression with its unique advantageous including fast onset of function, economic low-cost as well as minimal side effect [10, 11]. However, the involved pathway underlying the light effect on mood is not well understood, particularly little is known about the visual pathway with light therapy for non-seasonal depression [12].

As a key source of serotonin (5-HT), dorsal raphe nucleus (DRN), controls 5-HT release into the forebrain and has a vital role in mood regulation [13, 14]. Currently available antidepressants like selective serotonin reuptake inhibitors (SSRIs) mainly based on the modulation of raphe system to reduce 5-HT turnover [15, 16]. A mouse model with tryptophan hydroxylase-2 (TPH-2) mutation in raphe nucleus, which was analogous to human mutation originally identified in a late-life depression cohort [17], showed significant depression-like behavior with the functional 5-HT deficiency [18]. On the other hand, the retinal projection into DRN, namely retino-raphe projection, exists in many rodents and primates [19–21], including the cat [22], Sprague Dawley rat [21, 23], Mongolian gerbil [20, 21], as well as monkey Cebus apella [19]. Even in the human brainstem an intensive response with the light stimuli can be detected using functional magnetic resonance imaging (fMRI) [2, 24]. Our previous study had indicated that retino-raphe projection could modulate serotonergic tone in DRN and subsequent affective behavior as well [20, 25]. Therefore, it is very likely that the retino-raphe projection may play an important role during the light therapy.

A depressive model induced with a high dose of stress hormone (as corticosterone in rodents) has been widely used to study the relationship between non-seasonal depression and adult neurogenesis [26–30]. High level of corticosterone could induce depression-like responses in rodents mainly through related glucocorticoid receptors (GR) and corticotrophin releasing factor (CRF) [31–33], also probably involved the pathway of phosphorylated cyclic AMP response element binding protein (CREB) and arrestin regulation [26, 32]. In this study, we used a rat model of depression induced by corticosterone administration in order to test the role of retino-raphe projection in the light therapy to non-seasonal depression. Our results revealed that this model was suitable for the study of light therapy and the retino-raphe projection was shown to be underlying the effect of light therapy for non-seasonal depression.

## Methods and materials

### Animals

Adult male Sprague Dawley (SD) rats (Rattus norvegicus, 250-280 g) were used in this study. They were kept in a 12:12 light / dark cycle (light cycle with 30-40 lux; dark cycle with ~0 lux) with food and water provided *ad libitum* (lights onset at 7:00 am; offset at 7:00 pm). All experiments were performed in accordance with policies on the utilization of animals and humans in neuroscience research and approved by the Faulty Committee on the Use of Live Animals in Teaching and Research (CULATR) in The University of Hong Kong, and the counterpart in the Shenzhen Institute of Advanced Technology (SIAT), Chinese Academy of Sciences (CAS), China.

### Corticosterone and fluoxetine administration

Corticosterone (CORT, C2505; Sigma-Aldrich, USA) at a concentration of 40mg/kg dissolved in sesame oil (Sigma-Aldrich, USA) was injected into SD rats for 14 days, according to the method of our previous work [34, 35]. Briefly, a stock emulsion of CORT was prepared daily by vortexing corticosterone in sesame oil for 10 min, followed by 60 min of sonication. Prior to every injection, the emulsion was vortexed briefly and injections were made subcutaneously in the neck region of the rat every 24 h. Control treatment was the same as above, but seasame oil was injected without CORT. The administration of fluoxetine (F132, Sigma-Aldrich, USA) was conducted according to previous reports [25, 36, 37]. Some rats were undergone daily intraperitoneal (ip) injection of fluoxetine for 14 days with the daily CORT injection. Each rat received injection of 10 mg/kg/day fluoxetine dissolved in saline each afternoon, after subcutaneous injection of CORT.

### Bright light therapy

During the last 7 days of CORT injection, some rats in light-treated groups received the light therapy (n=6). Under the regular 12:12 L/D cycle of ambient light, blue (470 ± 10 nm) and full white light were respectively provided by light-emitting diodes (LED) placed approximately 30-50 cm above the rat cages. The intensity of LED light (100 ~ 400 lux) was detected by Lux meter (TM-209M, Tenmars, USA). The intensity of ambient light was only around 35 lux. While the intensity of blue light or white light was detected by lux meter at the bottom of the cage with the onset of ambient light, in order to match the real environment when they received the light therapy in morning. The light exposure was performed for 30 min each morning during the last 7 days of CORT injection. Onset of light therapy was at 7:00am each day, with the same onset time of ambient lighting. For an experiment of c-Fos test at the last day, those mice were respectively perfused with deep anesthesia after they were kept at the dark condition for 45 min following the 30 min duration of bright light therapy.

### Force swim test (FST)

Rats were placed in a cylindrical glass tank filled with water (21-22°C) to a depth of 30 - 40 cm and the whole test was recorded by video. On the first day, the animals were placed in the cylinder for a 15 minute pre-exposure. Data were collected during a second test conducted 24 hours later. Then the animals were monitored for a period of 5 minutes by trained observers. The analysis of animal behavior included three types of behavior in the test, namely climbing, swimming and immobility. The definition of behavioral types and more detailed procedure were described previously [35, 38].

### Sucrose preference test (SPT)

At the last four days of CORT or oil injection, the SPT experiment was started. The animals were firstly trained to drink a 1% sucrose solution by exposing them to sucrose solution instead of tap water for 24 h. They then were given a continuous 72 h two-bottle exposure to the 1% sucrose solution bottle and tap water bottle with the left/right location balanced across the cages. In order to prevent the possible deviation, two bottles were kept at same conditions using balanced location and were reversed the left/right location every night. At last 2 days, the bottles were weighed every 24 h at 10:00 am in the morning. The sucrose preference was determined as

(Δ weight of sucrose) / (Δ weight of sucrose +Δ weight of water) × 100 ^[39]^.

### Open field test (OFT)

The open field is a square arena (80cm length x 80cm wide x 40cm deep) with metal floors and wooden walls. Rats were placed in the center of the open field and allowed to freely explore the arena during a 10-minute test session. Both central and peripheral activities were measured using an automated video-tracking system. Percent of time in the center was defined as the percent of total time that rats spent at the central 26×26 cm area of the open field. Enhanced exploration of the unprotected central portion of a novel open field and reduced thigmotaxis were correlated with reduced anxiety levels [40].

### Corticosterone detection via ELISA

Blood samples of rats were collected at 19:00 pm, about 1 hour after the last CORT injection on the final day. Rats were anaesthetized and blood samples were collected from their tail vein within 3 min using a heparinized needle. Approximately 0.3 ml of blood was collected into a clean 1.5 ml microcentrifuge tube. Centrifugation (1000 rpm, 30 min and 4°C) of the samples was conducted then the serum was transferred into clean tubes and stored at −80°C. The CORT level was determined via a CorrelateEIA corticosterone kit (Assay design, USA). It was performed according to the manufacturer’s instruction and was described previously [28, 34, 35].

### Selective elimination or activation of retino-rapha circuitry

The method using CTB injection into DRN followed by intraocular injection of anti-CTB-saporin conjugate could specifically eliminate the retino-raphe projection, which was described previously [25]. Briefly, rats were anaesthetized with ip injection of a mixture of ketamine and xylazine (2:1, v/v). They were placed in a stereotaxic apparatus and craniotomy was performed. A Hamilton syringe with a 27G needle was inserted and stereotaxically positioned into the DRN regions, then CTB-conjugated Alexa Fluor 488 was deposited into DRN. The injection volume was 0.8 μl/animal with 0.2 μl sesame oil. The surgical area was cleansed and the incised tissues sutured. Three days after CTB injection, the animals were intraocularly injected with anti-CTB-Saporin (1μg saporin toxin /eye, anti-CTB: saporin = 1:1 mol ratio, Advanced Targeting Systems, Inc. San Diego, CA). Injection of anti-CTB-Saporin was also performed for each animal in the anesthetized condition. Eyelids were gently retracted with fingers to allow the eyeball to protrude. The injection was performed using a Hamilton syringe with a 35G needle into the posterior chamber of the eye and the needle was left in place for around 4 minutes. After around 3 weeks of the surgery, the DRN-projecting retinal ganglion cells (RGCs) were specifically eliminated in the retina of each animal. The control animals received DRN injection of CTB, and then intraocular injection of saline instead of anti-CTB-Saporin.

Using a similar method, intraocular injection of AAV8-Syn-HM3q-mCherry was performed into 2 eyes of those rats, meanwhile the AAV8-Syn-mCherry was injected into other rats as control group (n=5). After 3 weeks to express those virus, one cannula (200 mm in diameter, NA: 0.37, RWD, China) was implanted into mid-DRN region (AP: –8.10 mm, ML: – 0.00 mm and DV: –6.30 mm). The rats were then given 1-week recovery time before behavioral FST experiment began. They were i.p. injected with CNO (0.5 mg/kg BW, Clozapine N-oxide, C0832, Sigma) 1hr followed by 2^nd^ section of FST experiment. Saline or fluoxetine injection was also conducted as negative or positive controls (n = 5 rats each group).

### Tissue collection and preparation

Following deep anesthesia, rats were perfused with saline (pH 7.4) followed by 4% paraformaldehyde (PFA) in 0.01M phosphate buffer (PB: pH 7.4). Retinas were collected followed by post fixation with 4% PFA for 45 min. Brains were removed from the skulls and stored in the 4% PFA solution at 4°C overnight. The next day, brains were rinsed in 0.01M PB and then stored in 30% sucrose in 0.01M PB until they sank. The hindbrain including the mid-brain raphe complex was then serially sectioned at 30 µm intervals using a microtone. Brain sections were collected as six alternate sets of slices (with each set containing one section at 180 µm intervals throughout the hindbrain) and stored at −20°C in a cryoprotectant storage buffer (30% ethylene glycol, 30% sucrose dissolve in 0.01M PB; pH 7.4) until they were used to perform immunohistochemistry [41, 42].

### Immunohistochemistry

Tissues were rinsed in 0.01M PBS (pH 7.4) three times with 10 minutes each time, then incubated in blocking solution with 5% donkey serum in 0.1% Triton-X-100 of PBS for 1 h, followed by incubation of primary antibodies for 48 h at 4°C. The tissue sections were washed by 0.01M PBS (pH 7.4) three times with 10 minutes each time, and then incubated in matched secondary antibodies (1:500, Molecular Probes) for 2 h at room temperature, including Alexa Fluor donkey anti-rabbit 488nm-conjugated IgG, Alexa Fluor donkey anti-sheep 568nm-conjugated IgG, Alexa Fluor donkey anti-rabbit 647nm-conjugated IgG and so on. All sections were washed by 0.01M PBS (pH 7.4) three times with 10 minutes each time, finally cover-slipped by aqueous mounting medium (Dako Corp., Carpinteria, California, USA) [43, 44]. The primary antibodies included rabbit anti-c-Fos antibody (1:1000, PC38 (Ab-5), Calbiochem), rabbit anti-serotonin (5HT) antibody (1:1000, S5545, Sigma), sheep anti-tryptophan hydroxylase (TPH) antibody (1:1000, T8575, Sigma), rabbit anti-corticotropin releasing factor (CRF) antibody (1:500, ab8901, Abcam), rabbit anti-glutamate antibody (1:1000, G6642, Sigma), mouse anti-GAD67 antibody (1:1000, MAB 5406, Chemicon), mouse anti-GABA antibody (1:1000, A0310, Sigma), rabbit anti-5-HT3A antibody (1:200; ab13897, abcam), mouse anti-parvalbumin (PV) antibody (1:1000, P3088, Sigma) as well as rabbit anti-somatostatin (SOM) antibody (1:100, AB5494, Millipore). In addition, stain of anti-glutamate, anti-GAD67 and anti-SOM antibodies on brain sections sometimes used a tyramide signal amplification (TSA, T20934, Molecular probes) kit according to the manufacturer’s protocol [45].

### Immunoperoxidase staining

A serial sections including the midbrain raphe complex were used for immunoperoxidase staining. The primary antibodies included those against c-fos (rabbit anti-c-Fos polyclonal antibody, 1:3,000; PC38 (Ab-5), Calbiochem (EMD Chemicals)), tryptophan hydroxylase (TPH; sheep anti-TPH antibody, 1:10,000; T8575, Sigma), phospho-CREB (p-CREB; rabbit anti-phospho-CREB monoclonal antibody, 1:1000; #9198, Cell signaling Technology), GAD-67 (mouse anti-GAD67 antibody, 1:1000; MAB 5406, Chemicon) as well as 5-HT3A (rabbit anti-5HT3A receptor antibody, 1:200; ab13897, abcam). The procedure for double immunostaining of c-Fos and TPH was described here briefly. Tissue serial sections were washed by 0.01 M PBS in 12-well plate, twice 15min with shaking; then rinsed in 1% H2O2 in PBS for 15 min, followed by washing in PBS for 15 min and pre-incubation in 0.1% PBST for 1 h. Sections were then incubated overnight at room temperature (RT) with rabbit anti-c-Fos antibody (1:3,000) in 0.1% PBST. After about 16 h incubation, the tissue was washed twice in PBS, 15min each time with shaking; followed by incubation with a biotinylated goat anti-rabbit secondary antibody (1:200, E043201, Dako) in 0.01 M PBS for 90 min. Tissue was washed 2×15 min in PBS and then incubated in an avidin-biotin-peroxidase complex (Elite ABC reagent, 1:200; PK-6100, Vector Laboratories) in 0.01 M PBS for 90 min. Tissue was then washed 2×15 min in PBS, and incubated in a peroxidase chromogen substrate (Vector SG, SK4700, Vector Laboratories; diluted as the instruction) in 0.01 M PBS for 20 min. After the chromogen reaction, tissue sections were immediately washed in 0.01 M PBS twice, 15min each time with speedy shaking; then rinsed in 1% H_2_O_2_ for 5 min and washing in PBS for 15 min followed by 1 h pre-incubation of 0.1% PBST. Then, sections were incubated with sheep anti-TPH antibody (1:10,000) in 0.1% PBST overnight at RT. All subsequent steps were identical to those described above for the staining of c-Fos antibody, except for the secondary antibody and chromogen reaction steps; in which a biotinylated rabbit anti-sheep secondary antibody (BA-6000, 1:200; Vector Laboratories), and a peroxidase chromogen substrate (DAB substrate kit, SK-4100, Vector Laboratories; diluted as the instruction) in the distilled water for 5 min. Finally, sections were washed twice in PBS to stop the reaction, then mounted on microscope slides and mounted with coverslips after dehydrated treatment. The color reaction of the c-Fos immunostaining will be blue-black and localized to the nucleus, whereas TPH immunostaining was orange-brown and localized to the cytoplasm. The immunostaining process of p-CREB (1:1000), GAD-67 (1:2000) and anti-5HT3A (1:200) was similar to those described above [41, 46].

### DRN location and analysis

Serial brain sections of DRN (30 µm) were obtained from adult male SD rats. One sixth sections were immunostained with a rabbit anti-5HT antibody (S5545, Sigma) or a sheep anti-TPH antibody (T8575, Sigma), which can indicate the area of DRN filled with serotoninergic neurons. According to the different distances from bregma, subregions of DRN can be divided into the rostral portion of DRN (~from −6.92mmm to −7.64mm), middle portion of DRN (~from −7.73mm to −8.45 mm) and caudal portion of DRN (~from −8.54mm to −9.26mm). Moreover, the three subregions of DRN can be further divided into dorsal part of DRN (DRD), ventral part of DRN (DRV), ventrolateral part of DRN (DRVL), interfascicular part of DRN (DRI) and caudal part of DRN (DRC). Then MRN represent median raphe nucleus, which does not belong to DRN. The anatomy of DRN was described according to a classic rat stereotaxic atlas [47]. Based on that method, quantification of serotoninergic neurons was performed in different subregions of DRN [48].

### Data analysis

Data were shown as means ± SEM. The student T-test was used for comparisons between two groups, whereas one-way ANOVA was applied to compare three or more groups of data, followed by Newman-Keuls or Tukey multiple comparison test using the Prism 5.0 software. Tukey multiple comparison test was marked at those results when it was able to choose to conduct further statistics. A probability (P) value of <0.05 was considered at the statistically significant level while the <0.01 was at the remarkably significant level.

## Results

### Light therapy reversed the depressive-like behavior in the stressed rats caused by corticosterone administration

A rodent model of depression induced by the high level of stress hormone, corticosterone (CORT) with 40 mg/kg/day for 14 days, was able to be evaluated by forced swimming test (FST) (Fig 1 A), sucrose preference test (SPT) (Fig 1 B) as well as body weight change (Fig S1), respectively. And this type of depressive-like responses caused by high intake of CORT could be reversed by a typical SSRI drug, fluoxetine (10 mg/kg/day for 14 days, Fig 1 A and B). Moreover, our data in the FST clearly indicated that both white and blue light therapy were able to significantly reverse the depression-like responses in those stressed rats (Fig 1 C-E and Fig S2). The blue light therapy was demonstrated to exhibit a better efficacy compared with that of white light therapy at the same intensity (Fig 1 C and D), especially using the 400 lux blue light therapy (470 nm, 400 lux, 30 min each morning for 7 days; see Fig 1C, E and Movie S1; typically, the change of immobility time at Fig 1E and 2A, p<0.001, Blue-400 or Saline vs CORT groups). The sucrose preference in the saline-injected group was also enhanced significantly with the blue light therapy (p<0.001, Saline vs CORT groups) although they had already received the CTB injection into DRN before initiating the light therapy (Fig 2B). While revealed by open field test (OFT) (Fig 2 C), those stressed rats did not display obvious anxiety-like behavior, which was consistent with some previous reports [35, 49]. Neither 100 lux blue light or 100 lux white light exhibited sufficiently significant efficacy (Fig 1 C,D and Fig S2). The red light of 100 lux that belonged to the highest intensity of red light we could apply with our light source was unable to reverse the immobility time of those depressed rats significantly (Fig S2 G).

**Figure 1.**
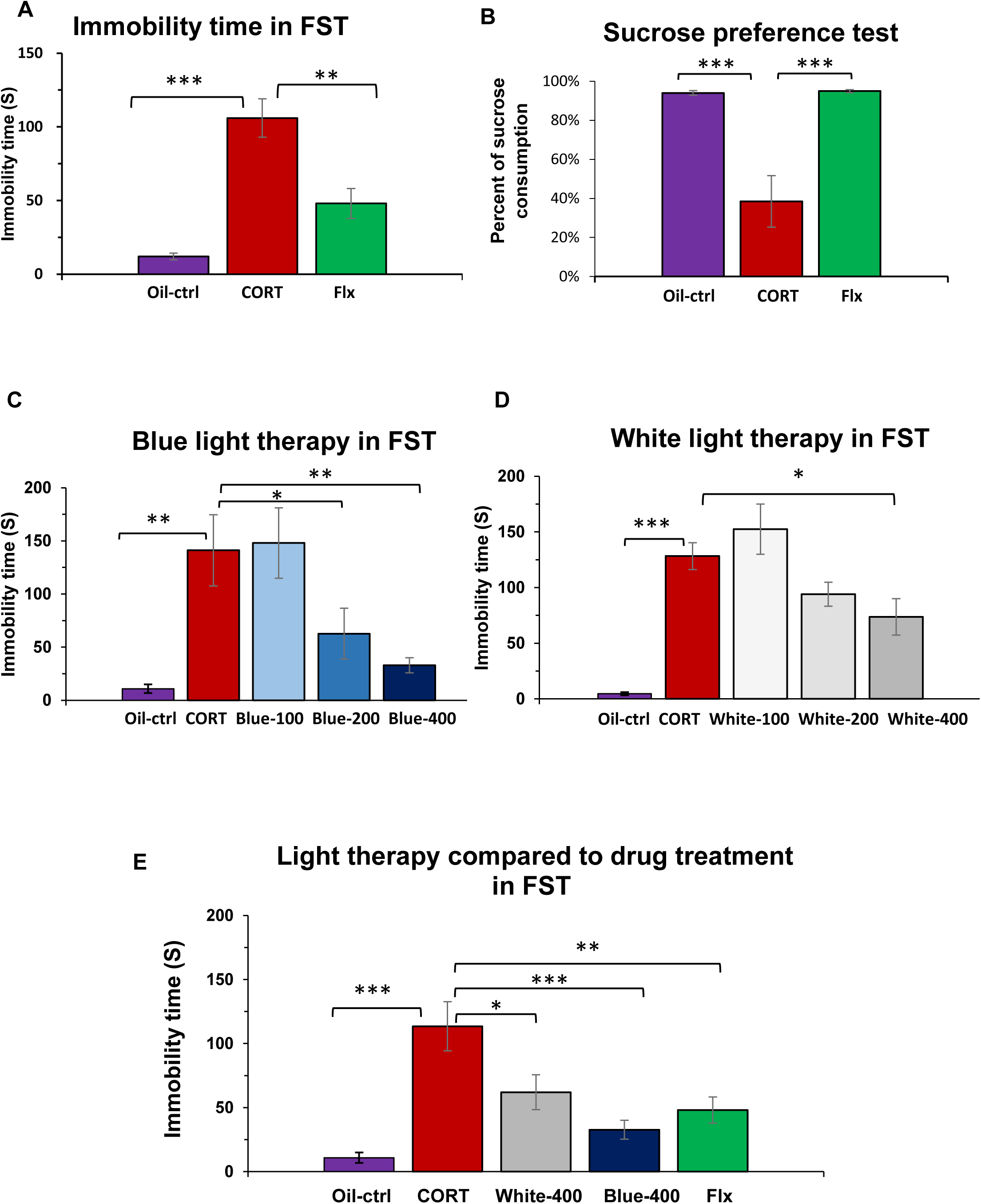
Light stimuli reversed the depressive behavior of stressed rats. (A) Immobility time of those stressed rats during FST. (B) Sucrose preference test of those stressed rats. (C) Blue light therapy at 100 lux, 200 lux and 400 lux. (D) White light therapy at 100 lux, 200 lux and 400 lux. (E) 400 lux light therapy compared with fluoxetine treatment. Note that the blue light therapy had better efficacy than the white light therapy at 400 lux, with similar to fluoxetine treatment. **\***P < 0.05, ** P < 0.01, ***p<0.001 vs. CORT group, n=6. Oil-ctrl: oil injected control group; CORT: corticosterone injected group; Blue-100, Blue-200 and Blue-400: light treated groups using blue light with 100 lux, 200 lux and 400 lux, respectively; White-100, White-200 and White-400: light treated groups using white light with 100 lux, 200 lux and 400 lux, respectively. Flx: corticosterone injection with fluoxetine treatment for 14 days.

**Figure 2.**
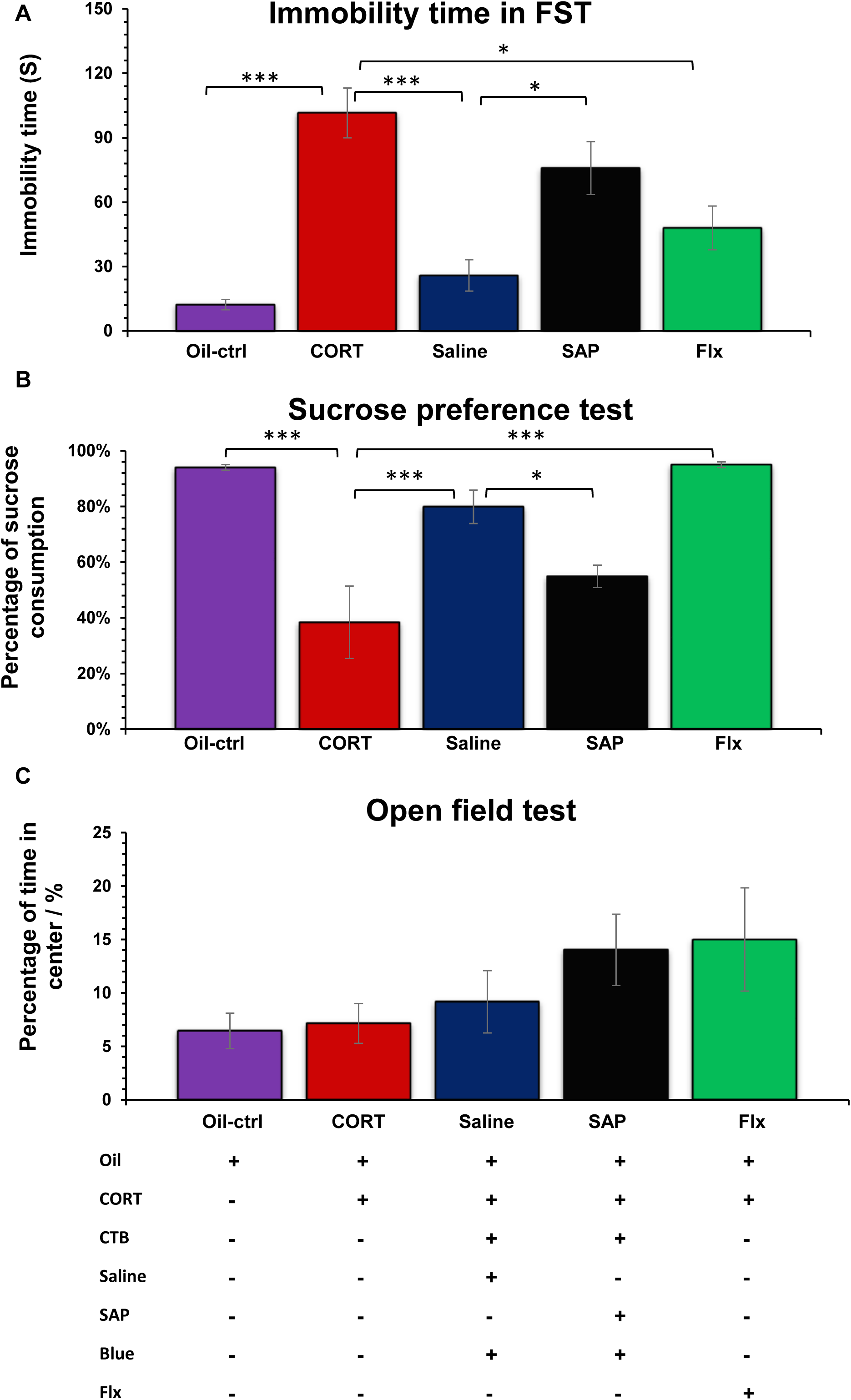
Eliminated retino-raphe projection diminished antidepressant effect of light therapy to stressed rats. (A) Forced swim test, (B) Sucrose preference test and (C) Open field test. Oil-ctrl: oil injection group, n=18; CORT: corticosterone injection group, n=24; Saline: CTB injection with saline injection, followed by corticosterone injection and blue light therapy (400 lux), n=8; Saporin: CTB injection with saporin injection, followed by corticosterone injection and blue light therapy (400 lux), n=8; Flx: corticosterone injection with fluoxetine treatment, n=6. * P < 0.05, ** P < 0.01, ***p<0.001 vs. CORT group.

Hence, it was indicated that the effect resulted from the light therapy for a week was sufficiently effective for those stressed rats to reverse their depressive behavior, in particular using the 400 lux blue light therapy. The rat retinas have been indicated to be very sensitive to the blue light (450 nm - 495 nm) through their classical photoreceptors and melanopsin photoreceptor [50, 51], which could be the main reason for the efficacy of blue light therapy over the white light therapy (Fig 1 E, p<0.001, Blue-400 vs CORT groups; p<0.05, White-400 vs CORT groups).

### Manipulation of retino-raphe circuitry effected on antidepressant function of light therapy

There was a distinct retino-raphe projection in the SD rat including average 700 DRN projecting RGCs each retina (Fig 3); among them, the majority belonged to alpha cells (approximately 72%, Fig 3C and S4) and around 10% melanopsin cells as well (Fig S4) (Li et al.). While Saporin (SAP) injection was performed to specifically eliminate their retino-raphe projection prior to the start of light therapy, the number of DRN projecting RGCs dramatically decreased from an average 700 to 150 cells in each retina (Fig 3, p<0.001, Saline vs SAP group, independent student’s t-test). Thus the antidepressant effect from the light therapy for those stressed rats has been significantly attenuated after elimination of their retino-raphe projection; this was strongly supported by behavioral tests including FST and SPT (Fig 2 and Fig S3). These results suggest retino-raphe projection mediates the effect of light therapy since the elimination of the retino-raphe projection led to the significant attenuation of the antidepressant effect resulted from the light therapy.

**Figure 3.**
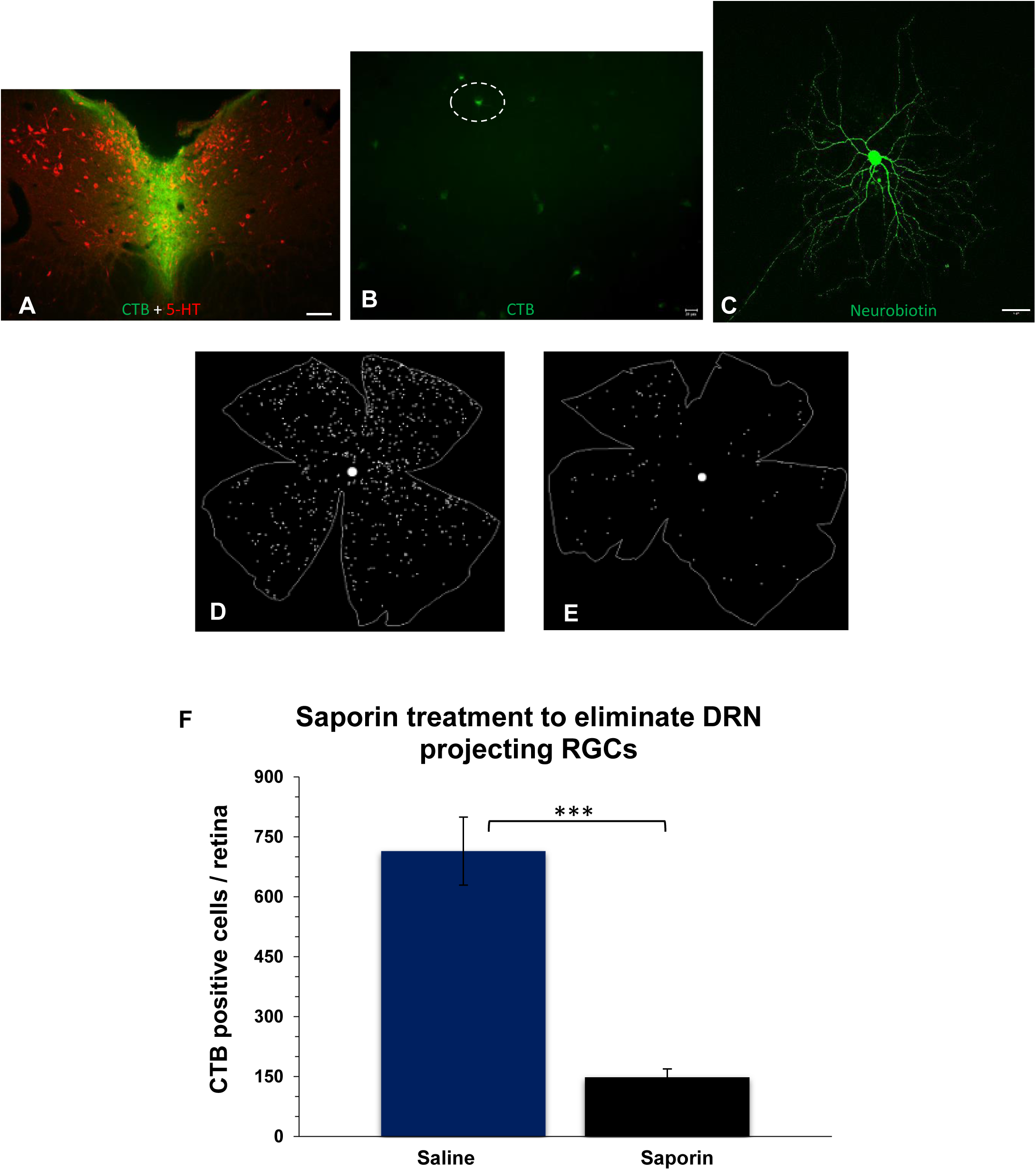
Saporin injection eliminated DRN projecting RGCs specifically. (A) is CTB injected site located at central DRN, (B) is CTB labelling cells on flat mounted retina and (C) is a CTB positive cell with intracellular injection. (D) indicates a distribution of CTB positive cells at saline-injected group (734 cells) while (E) is the saporin-injected group (79 cells). Note that (F) the number of DRN-projecting RGCs in the saporin injected group was significantly reduced compared with that in saline injected group. *** P < 0.001, n=6.

It was unable to completely abolish the DRN-projecting RGCs using those saporin treatment, but approximately 80% of DRN projecting RGCs was abolished (Fig 3), which was similar to some related studies [25, 52]. However, the attenuation of 80% retino-raphe connection was already sufficient to block the antidepressant effect from light therapy as both indicated by FST and SPT (Fig 2 A and B, p<0.001, Saline vs CORT groups; p<0.05, Saline vs SAP groups). It was not surprising that the stressed rats in the SAP-injected group still had a little increase of swimming time resulted from light therapy compared with that in the CORT group without the treatment (Fig S3 A). The reason was probably due to the ability of remaining DRN-projecting RGCs to still receive some signals from light therapy. Furthermore, the result of HM3q function to activate retino-raphe circuitry (Fig S3 C) consistently indicated the contribution of retino-raphe circuitry to antidepressant-like effect, also similar to the fluoxetine effect that was as positive control in this FST experiment.

### Significant c-Fos change happened in GABA cells in the subregion of DRN with corticosterone administration and light therapy

In terms of the c-Fos expression in the DRN (Fig 4 and Fig S5), those rats underwent noticeable alteration of c-Fos expression resulted from the high intake of stress hormone and also the effect of light therapy. DRN c-Fos expression was activated by corticorterone administration firstly then suppressed by light stimuli dramatically (Fig 4 and Fig S5), especially in the lateral wings of middle DRN, namely DRVL parts (Fig 4 F, p<0.05, Oil-ctrl vs CORT groups; p<0.01, CORT vs Co+Li groups; One-way ANOVA with Tukey’s multiple comparison test). Other subregions of DRN have not showed significant change between CORT group and light group except for the middle DRN and DRVL part (Fig 4 and Fig S5); this implicated the DRVL part of middle DRN was mainly charge of the noticeable change of c-Fos in whole DRN. A phosphorylated cyclic AMP response element binding protein (p-CREB) was regarded as the upstream factor of c-Fos [53, 54]; the trend of its changed supported the results of c-Fos alteration in DRN (Fig S6) although the p-CREB change was not statistically significant. Furthermore, as shown in Fig 4 G, the increase of c-Fos expression with corticorterone administration predominantly accounted for the activation of GAD67 positive GABA neurons (Fig 4 G, increased from 32% to 78% about GAD67 positive c-Fos cells; Fig S7 C, also increased from 15% to 55% about c-Fos positive GABAergic cells), rather than TPH positive serotonergic neurons (Fig S7 D and E, remaining ~25% TPH positive c-Fos cells with ~5% c-Fos positive TPH cells, no significant change). The hyperaction of GABAergic interneurons further induced the 5-HT synthesis inhibition indicated by quantification of TPH positive cells (Fig 5 H), which most likely associated with the depressive-like response as showed an increase of immobility time in FST and a decreased sucrose preference in SPT (Fig 1) [55]. However, light signals transmitted from the retino-raphe projection suppressed the c-Fos expression in the DRN at the light time with or without CORT administration (Fig 4, Fig S7 A and B); this was in agreement with the previous report by Fite et al [56]. Light stimuli caused the polarization and deactivation of some GABAergic interneurons, in which the activated level reduced from ~78% to ~57% on GAD positive c-Fos cells (Fig 4 H, ** P < 0.01, Co+Li vs CORT groups; One-way ANOVA with Tukey’s multiple comparison test); and also lowered to ~25% from ~55% on c-Fos positive GABA cells (Fig S7 C, ** P < 0.01, Co+Li vs CORT group; One-way ANOVA with Tukey’s multiple comparison test). The change of c-Fos expression in the both types of neurons in DRN (Fig 4), implicated the response from GABAergic interneurons was mainly contributed to the light effect from retino-raphe circuitry. This was consistent with that the DRN-projecting retinal fiber was embedded with many puncta of GAD67 positive GABA cells (Fig 5 A-C and Fig S8 A1-A5), rather than that of 5-HT neurons which also co-localized with touched retinal fibers (Fig 5 C and Fig S8 F).

**Figure 4.**
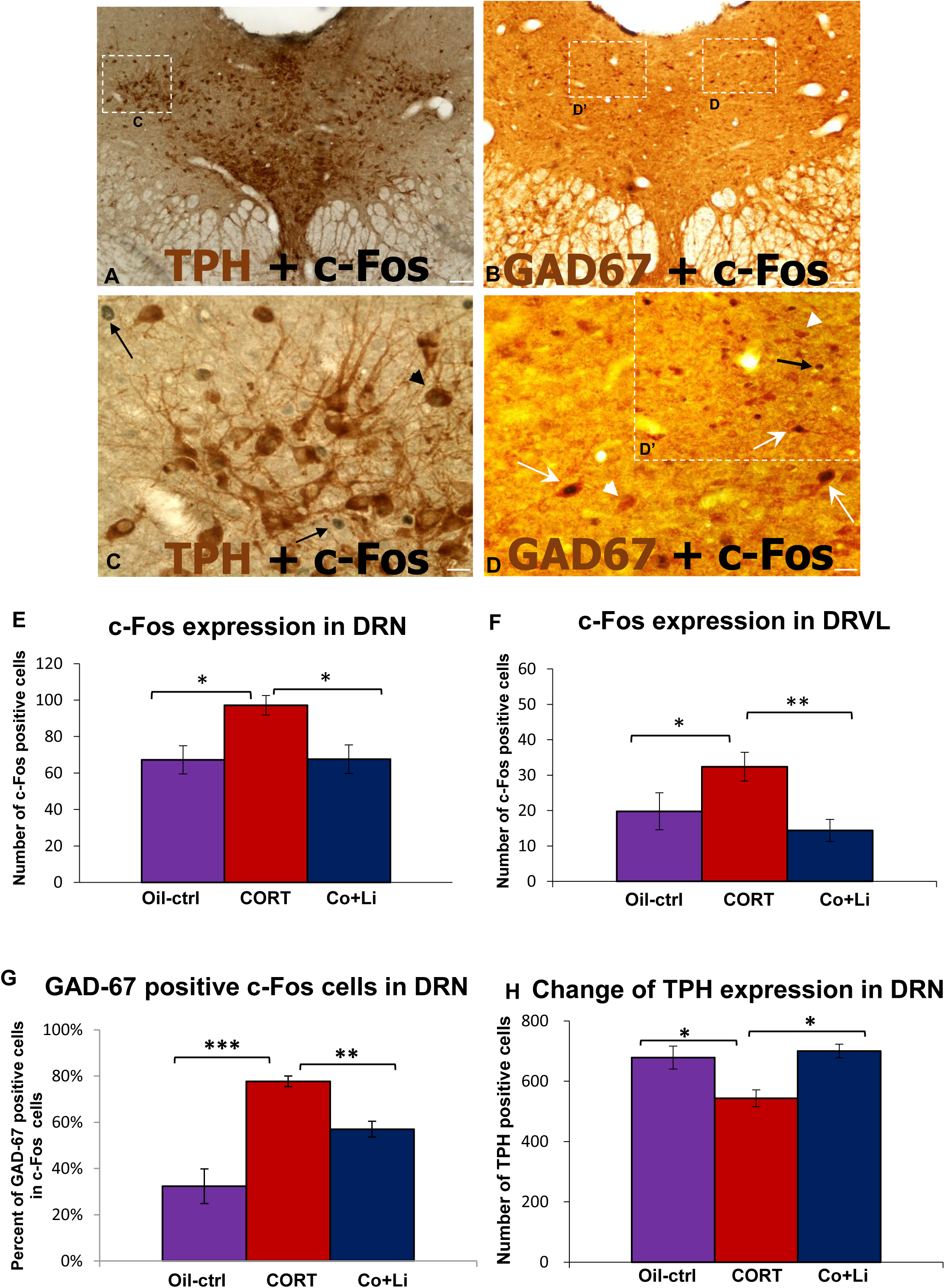
Altered expression of c-Fos in DRN with corticosterone administration and light therapy. Images (A) and (C) show the co-staining of c-Fos and TPH. Images (B) and (D & D’) indicate the co-stain of c-Fos and GAD-67 in DRN. Black arrows in (C) and (D’) show the c-Fos positive cells, while the black arrowhead at (C) indicates c-Fos co-labelling TPH cell; and the white arrowhead at (D) and (D’) indicates GAD-67 positive cell whereas the white arrows show the c-Fos co-labelling GAD-67 cells. Scar bars: A and B, 50 µm; C, D and D’, 10 µm. The bar graphs from (E)-(G) represent the change of c-Fos expression and (H) indicates the change of TPH positive cells by means ± SEM (n=6). Note that the different expression of c-Fos significantly occurred at GAD-67 positive GABA cells. *P < 0.05, ** P < 0.01, ***p<0.001 vs CORT group. Oil-ctrl: oil injected control group; CORT: corticosterone injected group; CO+Li: corticosterone injected with light treated group.

**Figure 5.**
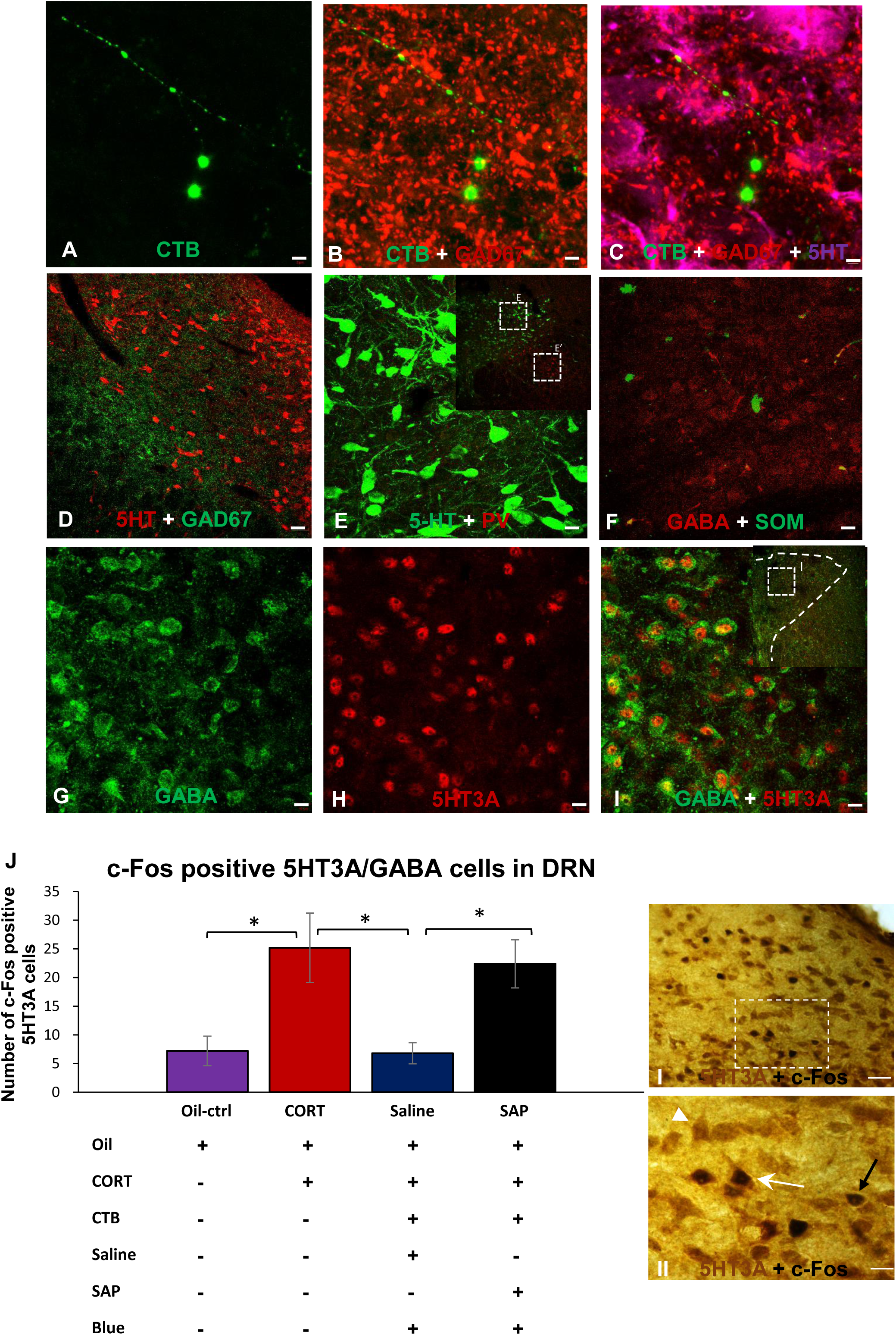
The hyperaction of 5-HT3A positive GABA cells was normalized by Light therapy. (A-C) indicate the triple labelling including CTB positive retinal fibers, GAD-67 positive GABA puncta and 5-HT positive cells. (D) GAD-67 positive GABA cells and their punta preferred to locate at the lateral parts of DRN (DRVL). (E & E’) and (F) The subtypes of GABA cells, parvalbumin (PV) positive GABA cells and somatostatin (SOM) positive GABA cells. Also the 5-HT3A positive GABA cells (G-I). Note that 5-HT3A positive cells, rather than PV cells and SOM cells, predominated the GABA expression in the DRN. The hyper activation of 5-HT3A positive GABA cells with c-Fos expression was induced by CORT administration, while it was normalized by light therapy in saline group (J). *P < 0.05 vs. CORT group, n=6. (I)-(II) presents the co-stain of 5-HT3A and c-Fos at (J). The black arrow indicates a c-Fos positive cell; and the white arrow indicates a c-Fos co-labelling 5HT3A positive cell whereas the white arrowhead shows the c-Fos negative 5HT3A positive cell. Oil-ctrl: oil injected control group; CORT: corticosterone injected group; CO+Li: corticosterone injected with light treated group. Scar bars: A-C, 5 µm; D, 50 µm; E-I, 20 µm; I, 20 µm; II, 10 µm.

### 5-HT3A coexpressing GABAergic interneurons in DRN mediated the function of light therapy for depression

The change of c-Fos was further supported by the microstructure of DRN in the rat. The distribution of GAD67 positive GABAergic neurons preferred to be located in both of ventrolateral parts of middle DRN (DRVL) (Fig 5 D and Fig S8 B), whereas the density of 5-HT neurons in DRVL was significantly lower than in the midline DRN (Fig 5 D). In agreement with previous study [21, 23], the retinal fibers from retino-raphe projection also innervated more in the DRVL part of the middle DRN (Fig 5 A and Fig S8 A). In addition, most of 5-HT cells expressed glutamate at the same time (Fig S8 D) [57], then 3 subtypes of GABA neurons could be observed in the rat DRN (Fig 5 and Fig S9), including parvalbumin (PV) positive GABA cells (Fig 5 E and Fig S9 D-G), somatostatin (SOM) positive GABA cells (Fig 5 F and Fig S9 H1-H3), and 5-HT3A positive GABA cells (Fig 5 G-I and Fig S9 A-C). More interestingly, the number of PV positive GABA cells was sparse in DRN and they preferred to locate at two pieces of medial longitudinal fasciculus (mlf) out of DRN (Fig 5 E and S9 D-G); and also the SOM expression was extremely weak in the DRN, even after using TSA amplification kit, only several cells with very weak signal were observed (Fig 5 F and Fig S9 H1-H3). Whereas 5-HT3A positive GABA cells distributed robustly in the whole DRN (Fig 5 G-I and Fig S9 A-C).

Thus we further detected the activation level of 5-HT3A positive GABA cells with the hormone stress and light treatment. The data indicated that 5HT3A positive GABA cells in DRN have been normalized the level of c-Fos expression with the light therapy (Fig 5 J, at II image, a white arrow indicates a c-Fos positive 5HT3A cell while a white arrowhead presents a c-Fos negative 5HT3A cell), which was initially activated by the high intake of stress hormone (Fig 5 J). While in the saporin-injected group the normalization of 5HT3A positive GABA cells was significantly blocked due to the disconnection of retinas and DRN (Fig 5 J, P < 0.05, Saline vs Saporin groups; One-way ANOVA with Tukey’s multiple comparison test). It was implicated that the level of c-Fos expression in 5-HT3A positive GABA cells was most likely to contribute to the light therapy effect through retino-raphe projection.

## Discussion

Depression is a serious mood disorder with a high prevalence and complicated pathophysiology [58, 59]. Currently available antidepressant drugs often had their limitations, like a long time lag for efficacy (several weeks or months), body weight gain and leading to sexual dysfunction [60, 61]. Nevertheless, to our knowledge, light profoundly effected on human mood state and cognitive function [1, 2, 62], and then the light therapy in clinical trials displayed its unique advantageous to treat depression, including fast-onset, low-cost and minor side effect [11]. However, the mechanism underlying them is still not clear so far. The present study revealed that the nerve innervation of retino-raphe projection had a key contribution to the light therapy for this non-seasonal depression model.

Our results at rodent model with corticosterone administration indicated light therapy especially the blue light therapy (200 ~ 400 lux) could effectively reverse the depression-like responses without seasonal pattern. The effect of light therapy was consistent with some clinical reports concerned the efficacy of light therapy, including the effect of light therapy usually needed a week, and the light therapy using blue light was often effective with lower light intensity compared with that using white light [12]. Before this study, several groups have already involved the animal study on light therapy for non-seasonal depression, not only for seasonal depression (SAD) [63, 64]. However, opposite results were obtained about light exposure to rodents [65, 66]. The light parameters and light conditions seemed to be crucial since an aberrant light condition could lead to impairment on mood [67]. This is suggested that it was more likely to benefit from the light therapy conducted in morning with the regular light/dark cycle [68]. A previous study has indicated that light signals could transmit into DRN directly in rats [56], and our previous work in gerbils has also revealed that light signals with photoreceptor ablation using MNU drug mainly driven by OFF alpha cells could modulate 5-HT production and then the related affective behavior through retino-raphe projection [25]. In this study, the blue light therapy was indicated to exhibit better efficacy than the white light therapy. Main reason was because DRN projecting RGCs in rats contained diverse types, at least 72% alpha cells and 10% melanopsin cells (Fig 3 and Fig 4S). Moreover, the rat retinas had been demonstrated to be very sensitive to the blue light (450 nm-495 nm) using their classical photoreceptors and melanopsin photoreceptor together [51]: the rat rods preferred to ~500 nm [50], and the rat cones were most sensitive at either 510 nm or 359 nm [69], whereas the intrinsically photosensitive retinal ganglion cells (ipRGCs, namely, melanopsin cells) in rats were most sensitive at 480 nm [51]. All of photoreceptors had some plain spectrum along their peak photosensitivity; then our using blue LED had the wavelength of 470 ± 10 nm belonged to the relatively much sensitive range of rat retina, which could reasonably activate most of retinal neurons especially with the little ambient white light (~35 lux). The 400 lux blue light therapy (470 nm, 400 lux, 30 min each morning for 7 days) had the best effect of light therapy for non-seasonal depression in this study, almost similar to the efficacy of fluoxetine treatment for 14 days (Fig 1, Fig 2 and Fig S3). So far there was no observational phototoxicity to be found in the retinal neurons following the stimulation of 400 lux blue light, and further evaluation is to conduct.

A recent report indicated that the projection of orexin neurons from the perifornical-lateral hypothalamic area (PF-LHA) into DRN displayed an important pathway in the light therapy, while using the Arvicanthis niloticus (Nile grass rat) as a model of seasonal depression with light deprivation [70]. In that case, the most reason might belong to that the grass rat is naturally lack of a direct connection between retinas and DRN region, unlike a distinct retino-raphe projection located at the SD rat [70, 71]. The immunotoxin induced elimination of the retino-raphe projection (lost 80% DRN projecting RGCs) significantly led to the attenuation of the antidepressant effect from the blue light therapy. This clearly revealed that the contribution from retino-raphe projection to the light therapy for those stressed rats. Another alterative argument was that after the retino-raphe projection was specifically eliminated, other retinofugal projections in the brain still probably participated in the light therapy. This could be the retino-SCN projection which was mainly driven by melanospin cells [72]. Since circadian phase shift due to altered light signal input into suprachiasmatic nucleus (SCN) was regarded as a main responsibility for light therapy to seasonal depression [73]. And the effect of circadian phase shift *via* photoentrainment was often taken with an advanced or delayed light stimuli [73, 74]. However, our light therapy conducted in those rats with regular light / dark cycle. Then other retinal projections could not be found to be associated with depression more than the retino-raphe projection. Therefore, our data in this study indicated that the rat retino-raphe projection was a key pathway for the function of light therapy to the non-seasonal depression caused by overdose stress hormone.

As an immediate early gene, *c-fos* with its protein (c-Fos) was often associated with cell depolarization [75, 76]. A dominant pathway for c-Fos induction involved an increased intracellular calcium or an increased levels of cyclic AMP. Previous reports have revealed that with the c-Fos expression, activation of GABAergic interneurons in DRN involved the depression-like responses of rodents [43, 77–79], since GABAergic interneurons in DRN particularly in DRVL part had a vitally inhibitory role in the modulation of 5-HT release in the raphe [14, 43, 80]. According to previous reports that the abundant expression of GR and CRF-2 receptors had been revealed in the DRN [81–83], suggesting that high intake of stress hormone could directly influence the DRN neurons with the hypothalamic–pituitary–adrenal (HPA) axis [83, 84]. Due to the high intake of corticosterone, CRF robustly expressed in DRN (Fig S8 E) and c-Fos was highly induced to express in the GAD67 positive GABA cells (Fig 4 and Fig S7); the inhibition of 5-HT neurons was then triggered (Fig 4 H) following the depressive-like responses happened in those stressed rats (Fig 1, 2 and 6) [31, 85]. When retinal fibers innervated into DRN with the input of light signals, they released glutamate to many types of neighbor cells [86–88], especially 5-HT cells and GABA cells in the DRN (Fig 5 and Fig S8). While the GABAergic interneurons received the glutamate from light signals could further communicate with 5-HT neurons probably through their receptors of GABA-A and GABA-B (Fig S10) [89–91]. The 5-HT neurons also might contact with the retinal fibers directly (Fig S8 A) [56]; and the vast majority of 5-HT cells belonged to glutamatergic (Fig S8 D) [57], to effect on neighbor GABAergic interneurons mainly through 5-HT2A, 5-HT2C as well as 5-HT3A receptors (Fig S10) [92–96]. However, c-Fos alteration in this study predominately occurred at GABAergic interneurons rather than 5-HT neurons (Fig 4 and Fig S7), which revealed the important role of GABAergic interneurons in mediating the effect of light therapy through retino-raphe projection. Further indication was that retinal fibers innervation, GABA cells distribution as well as c-Fos expression commonly preferred to exhibit at the DRVL part of middle DRN, where 5-HT neurons distributed relatively low (Fig 5 D and Fig S8 A) [97, 98].

There were mainly three subtypes of GABA cells located in the cortical and subcortical regions [99, 100], including PV positive cells, SOM positive cells as well as 5-HT3A positive cells. The 5-HT3A receptor, as the only ionotropic serotonergic receptor, had been revealed to express most, if not all, GABAergic interneurons that do not express PV or SOM protein at the mouse cortex [101]. In this study, GABAergic interneurons in the DRN were also found to mostly express 5-HT3A receptor (Fig 5 and S9). Compared with 5-HT3A positive GABA cells distributed robustly in the whole DRN, the population of PV positive GABA cells was sparse (Fig 5 E and Fig S9 D-G), consistent with the previous study [77]; and the SOM protein was only weakly expressed at a few GABA cells in DRN (Fig 5 F and Fig S9 H1-H) although previous reports had revealed its location in DRN [98, 102]. Thus we further detected the activation level of 5-HT3A positive GABA cells with the corticosterone administration and light therapy. Interestingly, it was found that light therapy has normalized the hyperaction of 5-HT3A positive GABA cells initially caused by high intake of stress hormone (Fig 5 J). When the retino-raphe projection was eliminated by Saporin treatment prior to light therapy, the normalization of the 5-HT3A positive GABA cells was lost, followed by the failure of light therapy to those stressed rats. It was further indicated that the 5-HT3A coexpressing GABA cells in the DRN mainly mediated the function of light therapy for non-seasonal depression through retino-raphe projection. Although PV positive GABA cells in DRN was also reported to involve the modulation of depression-like response in other type of rodent models [77], here 5-HT3A positive GABA cells were the key contribution to modulate the light therapy effect. In addition, 5-HT3A receptor activation and deactivation with c-Fos levels seemed to respectively associate with depression-like and antidepressant-like responses, which was in agreement with the function of 5-HT3A agonist and antagonist according to some previous reports [103–105].

In this study, our present data indicated that light therapy had a relatively rapid antidepressant effect at a week, compared with that antidepressant fluoxetine with the function onset after 2 weeks [58, 61]; thus this was less likely to associate with modulation of adult neurogenesis that usually resulted from chronic antidepressant effect [61]. The light stimulation did not significantly induce c-Fos immunoreactivity in the 5-HT neurons, but significantly deactivated the hyperaction of GABAergic interneurons and thus modulated the microcircuitry of DRN serotoninergic system. Light suppression of c-Fos expression was also reported previously but the molecular mechanism was not clear so far [56, 106]. C-Fos expression in glutamatergic neurons, like in the retina was usually implicated a maintenance of synaptic connection and signal transmission as well [107, 108], whereas GABAergic interneurons with c-Fos expression would be more likely to contribute an inhibitory function to their neighboring neurons [79, 80, 92, 93]. Before the light signals transmitted into the DRN, many GABAergic interneurons have already stayed in the activation state caused by high level of stress hormone, which was similar to the situation in DRN induced by other types of stress stimuli as well as the state of active sleep (namely, rapid eye movement sleep) [43, 77, 109]. The light signals through retino-raphe projection went into the DRN with the light stimuli and that was very likely to activate the arousal response, since the arousal response was a primary function of the DRN serotoninergic system [110, 111]; and the light stimuli could initiate arousal response rapidly [2, 112, 113]. While Saporin immunotoxin specifically eliminated the retino-raphe projection [25], the function of light therapy was significantly blocked. Thus the light therapy for non-seasonal depression is more likely to benefit from the light-induced arousal response in DRN serotoninergic system through retino-raphe projection. The function of retino-raphe projection with a natural optogenetics originated from retinal opsins [93, 114, 115], is not only for non-seasonal depression but might also contribute to seasonal depression and other circuitry-related affective disorders due to the intimate connection between raphe and other emotional areas [116–118], and seems to enable most healthy people to benefit from sunshine as well [119, 120]. In future the involved mechanisms on light therapy will be elucidated more thoroughly, which might benefit to understand the depression profoundly and well develop an advanced antidepressant way.

## Conclusion

This study indicated the retino-raphe projection had a function to modulate the neural activity in the DRN and contributed to light therapy for a non-seasonal depression model. The light stimulation especially using blue light could reverse the depression-like responses in those stressed rats caused by corticosterone administration. While the retino-raphe projection was largely eliminated using the Saporin immunotoxin, the effect of light therapy was attenuated significantly. During the light therapy for those depressed animals, the light signals through retino-raphe projection deactivated the hyperaction of 5-HT3A positive GABA cells with c-Fos expression that initially resulted from high intake of stress hormone; that eventually contributed to antidepressant effect of light therapy.

## Acknowledgments

This work was supported by Programme of Introducing Talents of Discipline to Universities (B14036), and Project of International, as well as Hong Kong, Macao & Taiwan Science and Technology Cooperation Innovation Platform in Universities in Guangdong Province (2013gjhz0002). This was also supported by grants from the Commission on Innovation and Technology in Shenzhen Municipality of China (JCYJ20150630114942262), the Postdoctoral Science Foundation of China (2015M582440), International Postdoctoral Exchange Fellowship Program 2016 by the Office of China Postdoctoral Council (20160021), and the National Key R&D Program of China (2017YFC1310503).

