## Supplementary material for "Light stimulation into dorsal raphe nucleus contributes to antidepressant effect for a stressed rat model": Legend S1-S10

**Supplementary data**

**Fig S-1 Rat model of CORT-induced depression.** Their behavior can be reversed by fluoxetine in forced swim test (FST, A-C) and sucrose preference test (SPT, D), and the change of body weight and glucocorticoids level in blood with corticosterone (CORT) administration (E and F). ***** P < 0.05, ** P < 0.01, *** p<0.001 vs. CORT group. Bar graphs represent the means$\pm$SEM (n=6). Oil-ctrl: control group with sesame oil injection; CORT: corticosterone injected group; Flx: corticosterone injected with fluoxetine treated group.

**Fig S-2 Blue light and white light treated those stressed rats with CORT administration.** At the same light intensity, blue light therapy had better efficacy than white light therapy, especially the 400 lux blue light therapy. *****P < 0.05, ** P < 0.01 vs. CORT group. Oil-ctrl: oil injected control group; CORT: corticosterone injected group; Blue-100, Blue-200 and Blue-400: light treated groups using blue light with 100 lux, 200 lux and 400 lux, respectively; White-100, White-200 and White-400: light treated groups using white light with 100 lux, 200 lux and 400 lux, respectively. Flx: corticosterone injection with fluoxetine treatment for 14 days.

**Fig S-3 Eliminated or activated retino-raphe projection effected on antidepressant function of light therapy in the CORT injected rats.** (A) Swimming time in forced swim test, (B) Climbing time in forced swim test. Oil-ctrl: oil injection group, n=18; CORT: corticosterone injection group, n=24; Saline: CTB injection with saline injection, followed by corticosterone injection and blue light therapy, n=8; Saporin: CTB injection with saporin injection, followed by corticosterone injection and blue light therapy, n=8; Flx: corticosterone injection with fluoxetine treatment, n=6. * P < 0.05, ** P < 0.01, ***p<0.001 vs. CORT group. (C) CNO effect on HM3q function to retino-raphe circuitry, while saline and fluoxetine injection were as negative and positive groups, respectively. N=5. * P < 0.05, ** P < 0.01 vs. saline group.

**Fig S-4 The types of DRN-projecting RGCs in the rat.** Bar graphs represent the means ± SEM. Percentage of alpha cells and other types RGCs mainly based on analysis of their cell morphology with the intracellular injection (n=50 cells, 4 rats). While ipRGCs were further identified by antibody (PA1-780) immunohistochemistry (n=6 retinas). ipRGCs: intrinsically photosensitive retinal ganglion cells.

**Fig S-5 Comparison of c-Fos expression in different subregions of DRN between the three groups.** Images of I-III indicate the subregions of DRN, which was defined by the expressed distribution of tryptophan hydroxylase (TPH). Bar graphs represent the number change of c-Fos expression using means$\pm$SEM (n=6). *P < 0.05, ** P < 0.01 vs. CORT group; One-way ANOVA with Tukey's multiple comparison test. Oil-ctrl: oil injected control group; CORT: corticosterone injected group; CO+Li: corticosterone injected with blue light treated group.

**Fig S-6 Comparison of p-CREB expression in DRN of different groups.** The mage indicates the high expression of p-CREB in the DRN with the outline of TPH co-stain. Bar graphs represent the number change of p-CREB expression using means$\pm$SEM (n=6), using the one-way ANOVA with Tukey's multiple comparison test. Oil-ctrl: oil injected control group; CORT: corticosterone injected group; CO+Li: corticosterone injected with blue light treated group. p-CREB: phosphorylated cyclic AMP response element binding protein; TPH: tryptophan hydroxylase.

**Fig S-7 Altered expression of c-Fos in GABA cells and 5-HT cells in DRN.** (A) and (B) show the c-Fos suppression of light stimuli without or with corticosterone administration. (C) and (D) indicate the c-Fos positive GABA cells and 5-HT cells in DRN, respectively. The bar graphs represent the means ± SEM (n=6). *P < 0.05, ** P < 0.01, ***p<0.001 vs CORT group; One-way ANOVA with Tukey's multiple comparison test. Oil-ctrl: oil injected control group; Oi+Li: oil injected with light treated group; CORT: corticosterone injected group; CO+Li: corticosterone injected with light treated group.

**Fig S-8 Retinal fibers projected into DRN and approached many types of cells.** (A) A retinal fiber (green) was embedded with many puncta from GAD67 positive GABA cells (red). (F1-F5) were serial images for (F) part with the interval of 150 nm, in which the arrows indicate the probably close interaction between the retina projecting fiber (green) and GAD67 positive puncta of GABA cells (red). (B) A GAD67 positive GABA cell body with many GAD67 positive puncta. (C) 5-HT cells (green) interacted with the puncta of GAD67 positive GABA cells (red). (D) Co-stain of glutamate and TPH antibodies, indicated most 5-HT cells expressed glutamate. (E) CRF expressed around the TPH positive 5-HT cells in DRN with CORT administration. (F) Retinal fibers (green) went through the 5-HT cells (purple). GAD67: glutamic acid decarboxylase-67, a key enzyme in GABA synthesis; TPH: tryptophan hydroxylase, a key enzyme in 5-HT synthesis; CRF: corticotropin-releasing factor. Scar bars: A and B, 10 µm; C, F and F1-F5, 5 µm; D, 20 µm; E, 40 µm.

**Fig S-9 Subtypes of GABAergic interneurons located in the DRN.** (A-C) indicate a subtype of GABA cells coexpressing 5-HT3A receptor robustly distributed in DRN. (D-F) show PV positive GABA cells located in DRN, but the number was few, only found two already marked by a circle. (G-I) present third subtype of GABA cells with SOM positive, whereas the signal was too weak to stain even using TSA kit to amplify when only several positive cells were observed with the weak expression of SOM. 5-HT3A: serotonin 5-hydroxytryptamine 3A receptor; PV: parvalbumin; SOM: somatostatin. Scar bars: A and C, 5 µm; B, D1 and E1, 50 µm; D2, D3 and E2, 20 µm; E3, 10 µm.

**Figure S-10 Main microcircuitry of DRN function involved the effect of light therapy to depressive rats.** High level of corticosterone intake could directly enhance the neuronal activity in DRN neurons especially the GABAergic interneurons in the DRVL part, mainly through related GR receptor and CRF factor, followed by the depression-like responses observed in rodents. Then the GABAergic interneurons co-expressing 5-HT3A received the light signals with glutamatergic input through retino-raphe projection and that could deactivate their hyperaction caused by corticosterone administration. They further communicated with 5-HT neurons mainly through their receptors of GABA-A and GABA-B, to modulate the 5-HT synthesis and release. The feedback from 5-HT neurons to GABAergic interneurons might act through 5-HT receptors such as 5-HT3A and 5-HT2C. And the direct effect of light signals on 5-HT cells through retino-raphe projection was not detected in this study, maybe only miner function. However, the depression-like responses in rodents were eventually reversed by light therapy through retino-raphe projection, with the 5HT3A positive GABAergic interneurons. CORT: corticosterone; GR: glucocorticoid receptors; CRF: corticotrophin releasing factor; GABA-A & GABA-B: 2 subtypes of GABA receptors; GLUT: glutamate; 5-HT3A, 5-HT2C: 2 subtypes of 5-HT receptors expressed at GABA cells, while only 5-HT3A was the ionotropic serotonergic receptor.
