## Supplementary material for "Light stimulation into dorsal raphe nucleus contributes to antidepressant effect for a stressed rat model": Fig S1-S10

### Figure S1

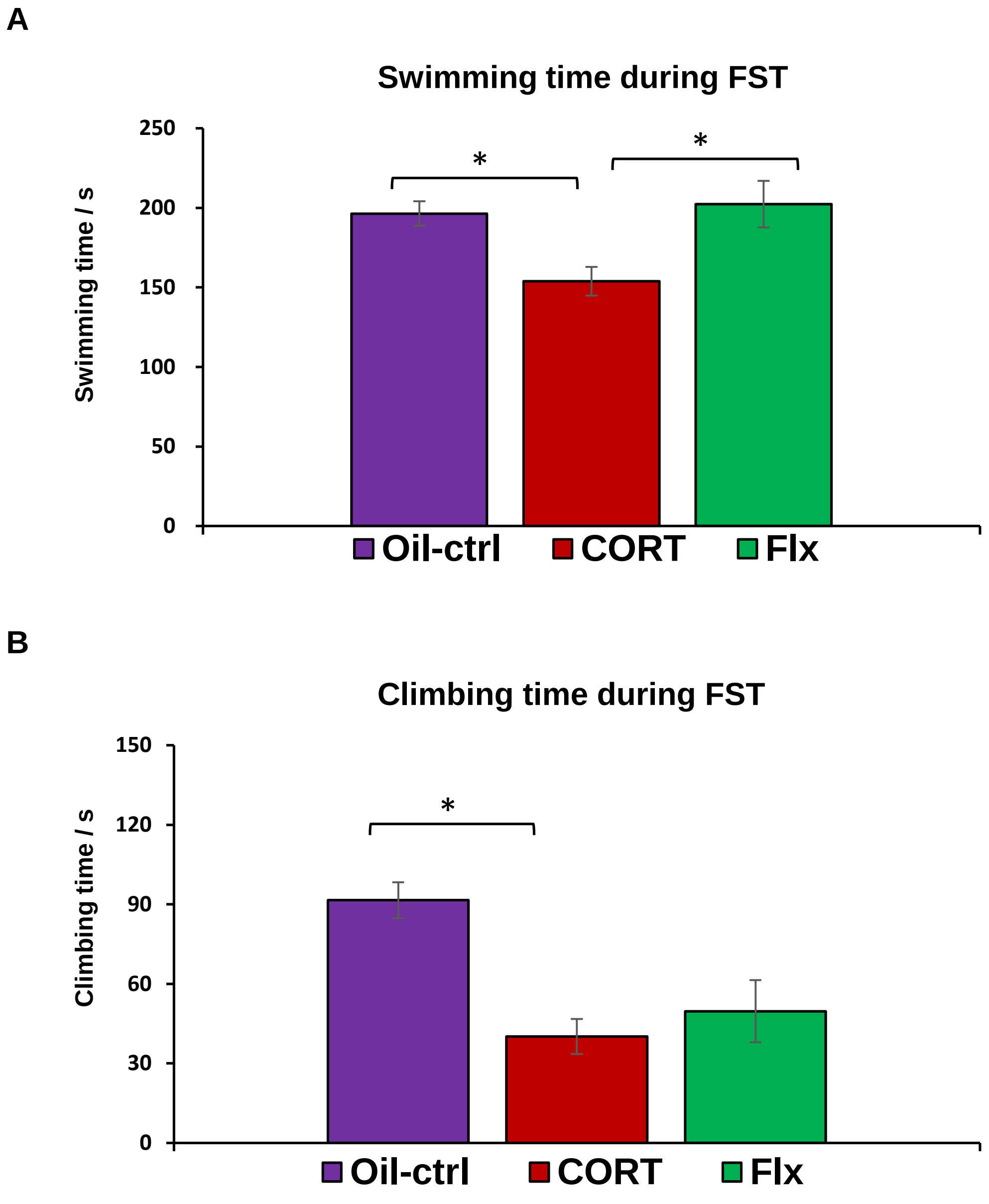

A
Swimming time during FST
Swimming time / s
250
200
150
100
50
0
*
*
Oil-ctrl
CORT
Flx
B
Climbing time during FST
Climbing time / s
150
120
90
60
30
0
*
Oil-ctrl
CORT
Flx

### Figure S1

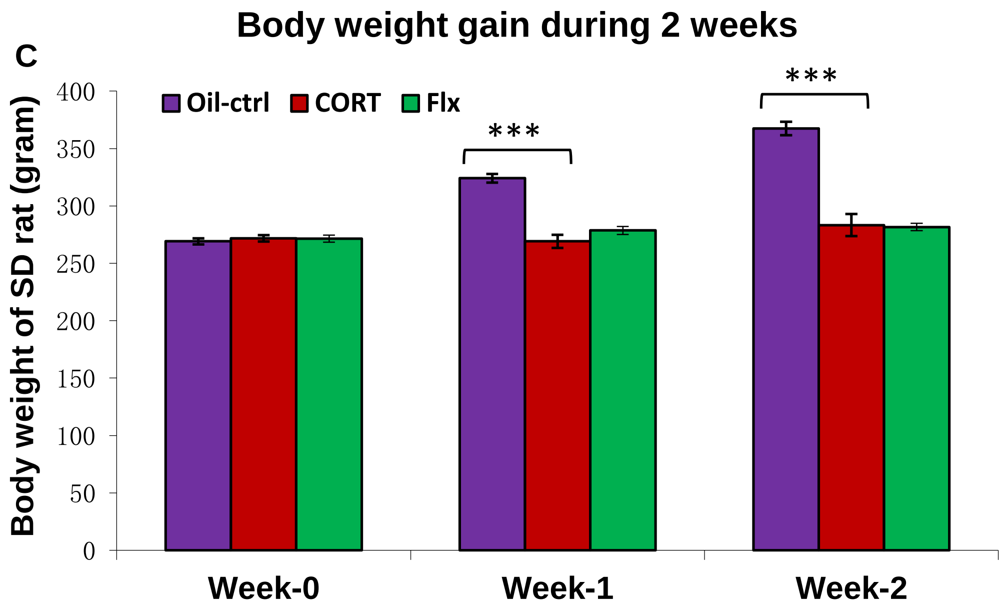
C
Body weight gain during 2 weeks
Body weight of SD rat (gram)
400
350
300
250
200
150
100
50
0
Oil-ctrl
CORT
Flx
***
***
Week-0
Week-1
Week-2

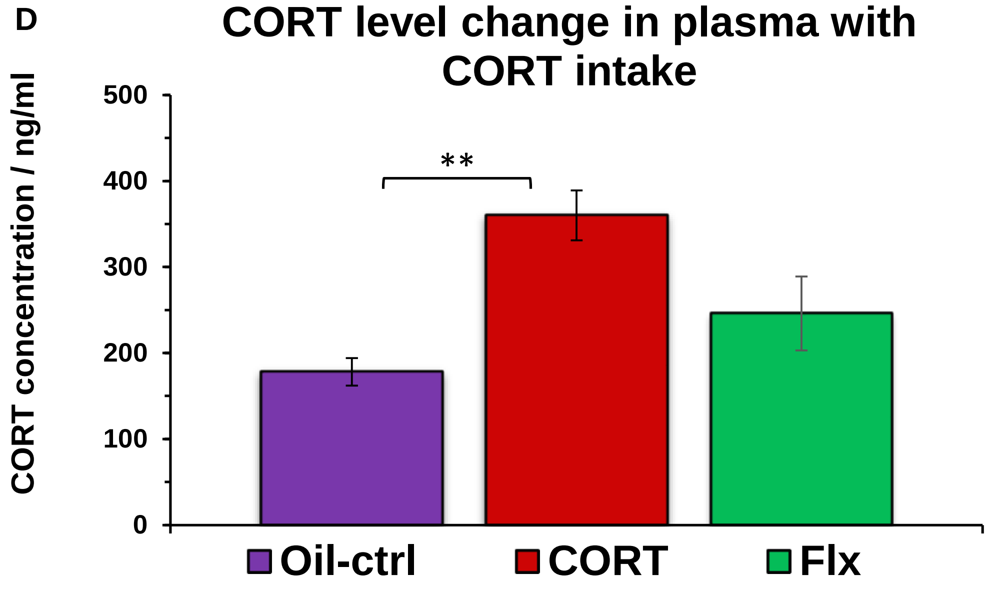
D
CORT level change in plasma with CORT intake
CORT concentration / ng/ml
500
400
300
200
100
0
**
Oil-ctrl
CORT
Flx

### Figure S2

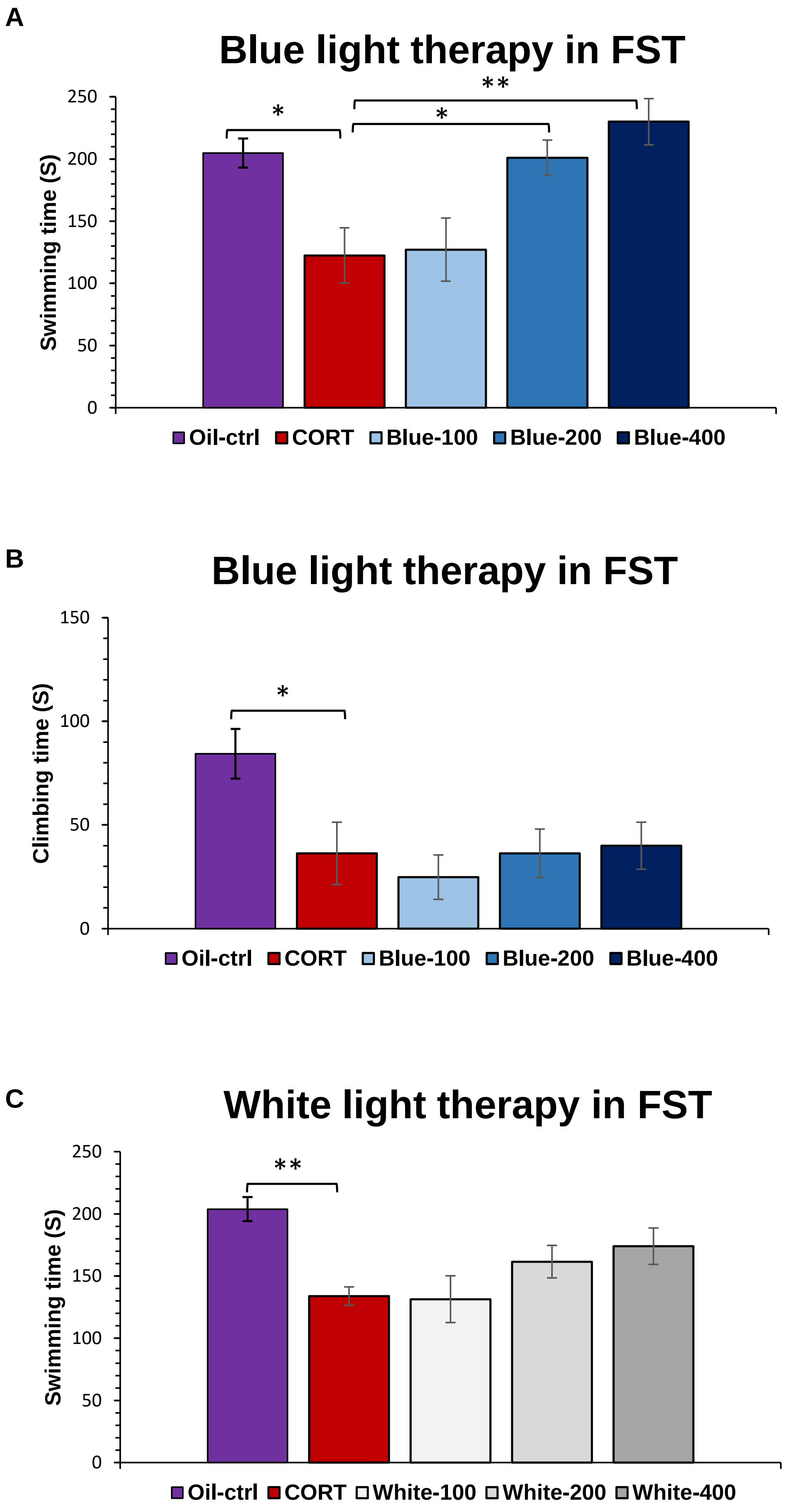

A
Blue light therapy in FST
Swimming time (S)
0
50
100
150
200
250
*
*
**
Oil-ctrl
CORT
Blue-100
Blue-200
Blue-400
B
Blue light therapy in FST
Climbing time (S)
0
50
100
150
*
Oil-ctrl
CORT
Blue-100
Blue-200
Blue-400
C
White light therapy in FST
Swimming time (S)
0
50
100
150
200
250
**
Oil-ctrl
CORT
White-100
White-200
White-400

### Figure S2

D

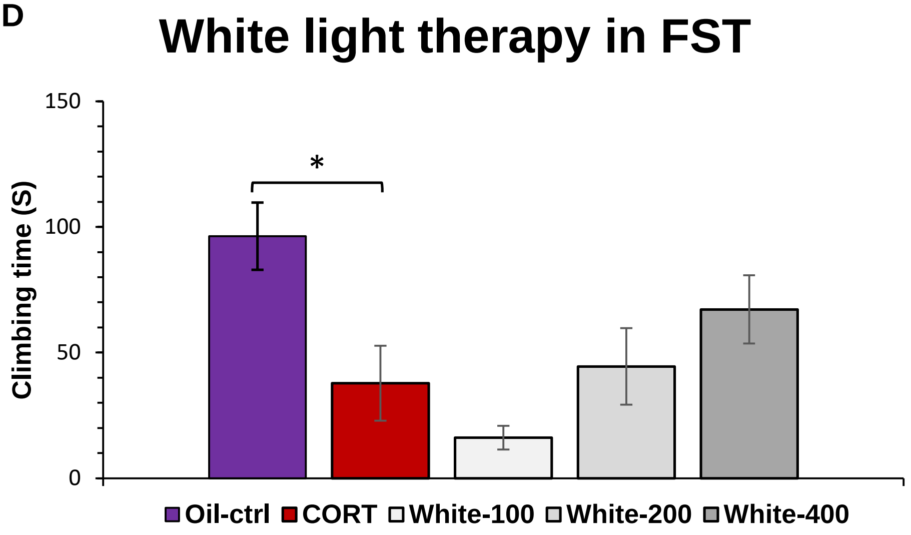


E

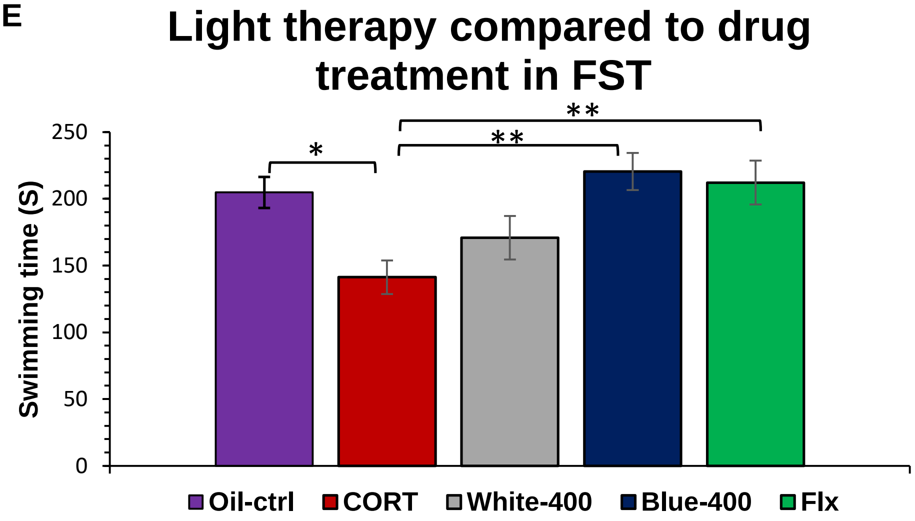


F

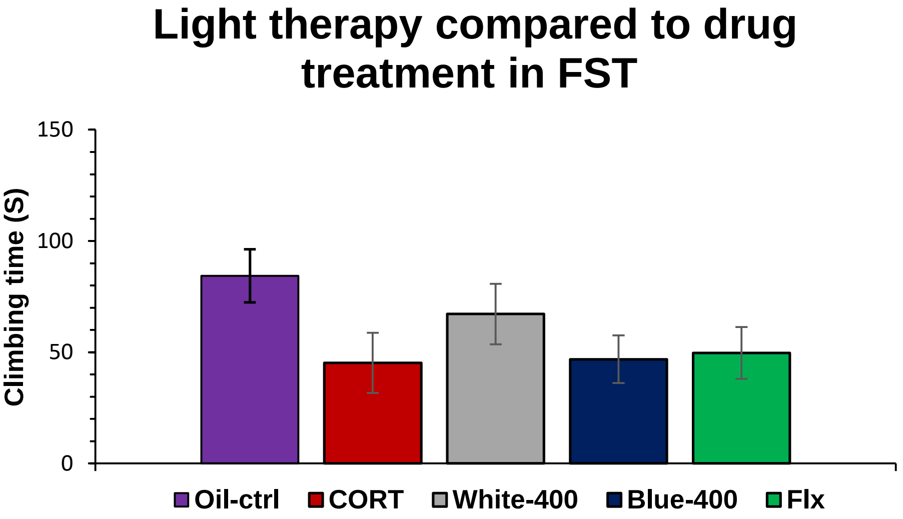

### Figure S2

G

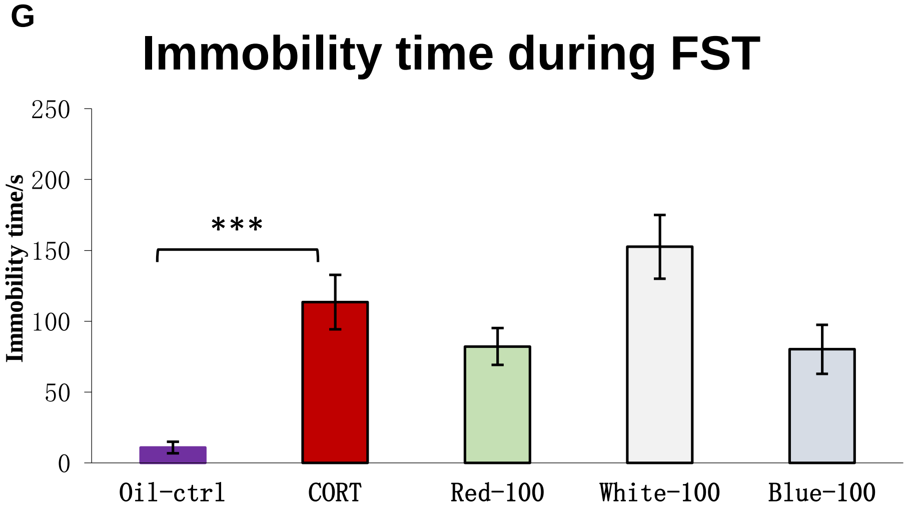


H

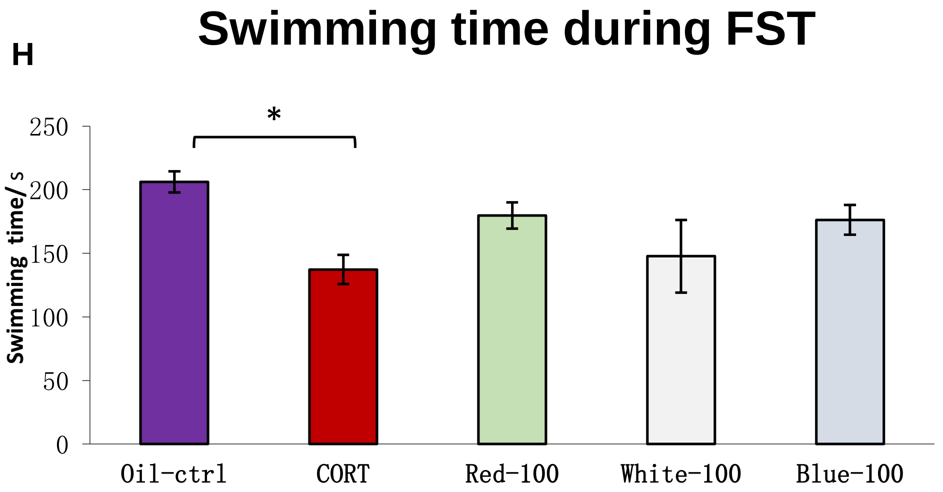


I

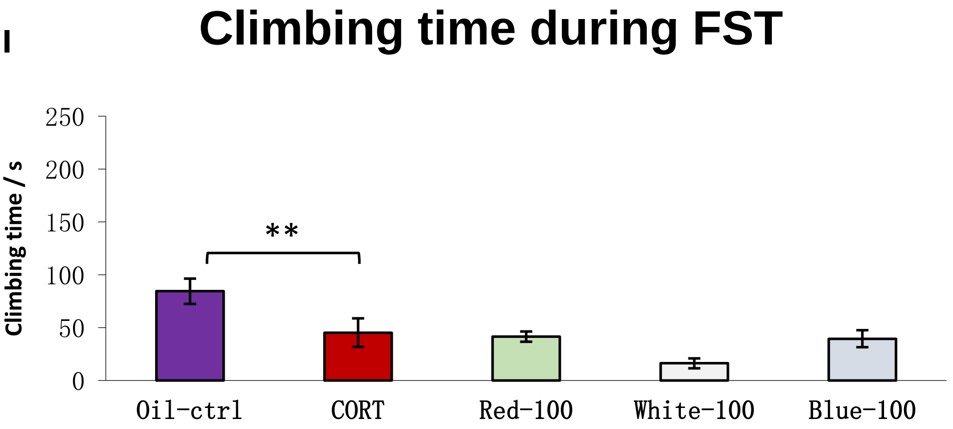

### Figure S3

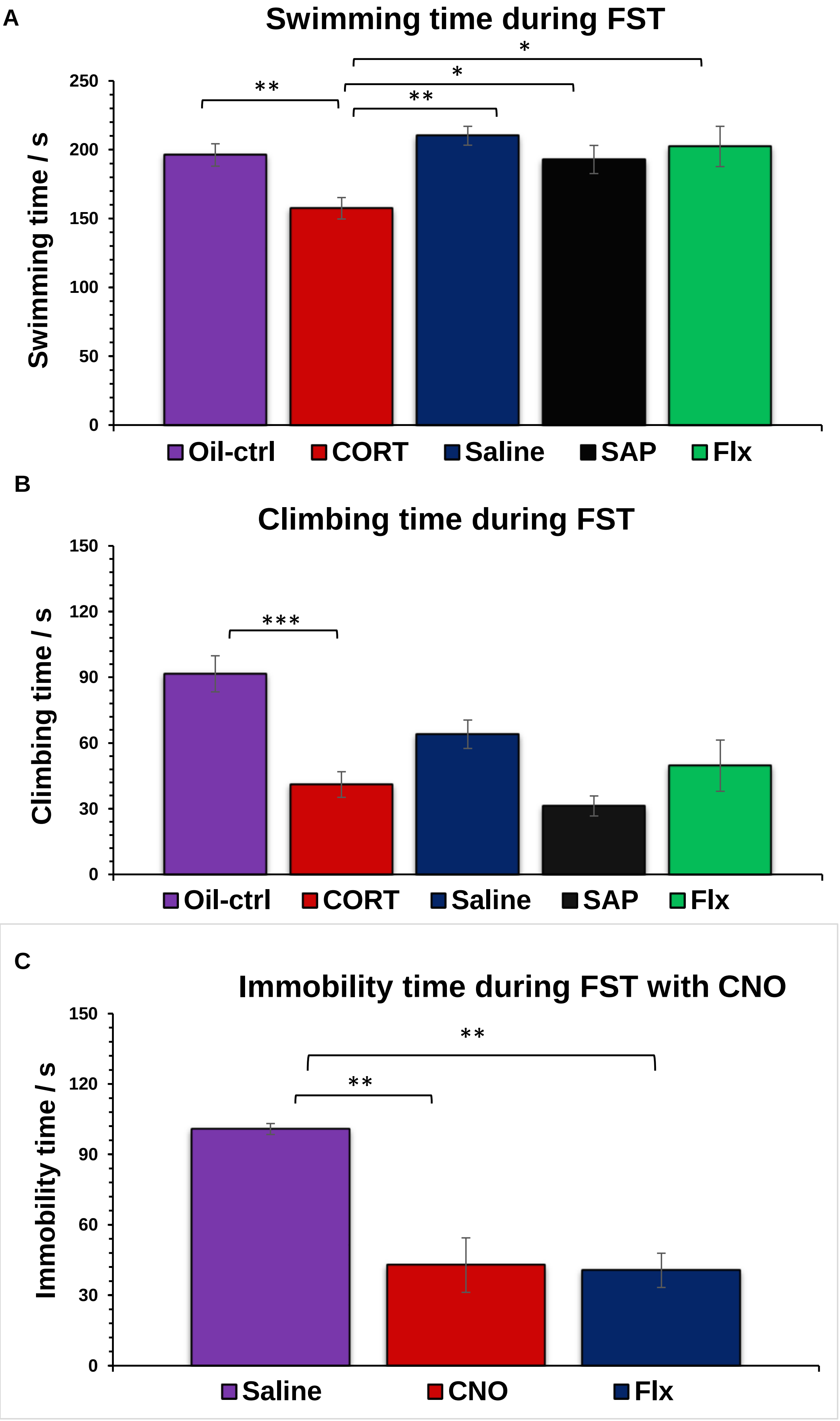
A
Swimming time during FST
Swimming time / s
250
200
150
100
50
0
**
*
*
**
Oil-ctrl
CORT
Saline
SAP
Flx
B
Climbing time during FST
Climbing time / s
150
120
90
60
30
0
***
Oil-ctrl
CORT
Saline
SAP
Flx
C
Immobility time during FST with CNO
Immobility time / s
150
120
90
60
30
0
**
**
Saline
CNO
Flx

### Figure S4

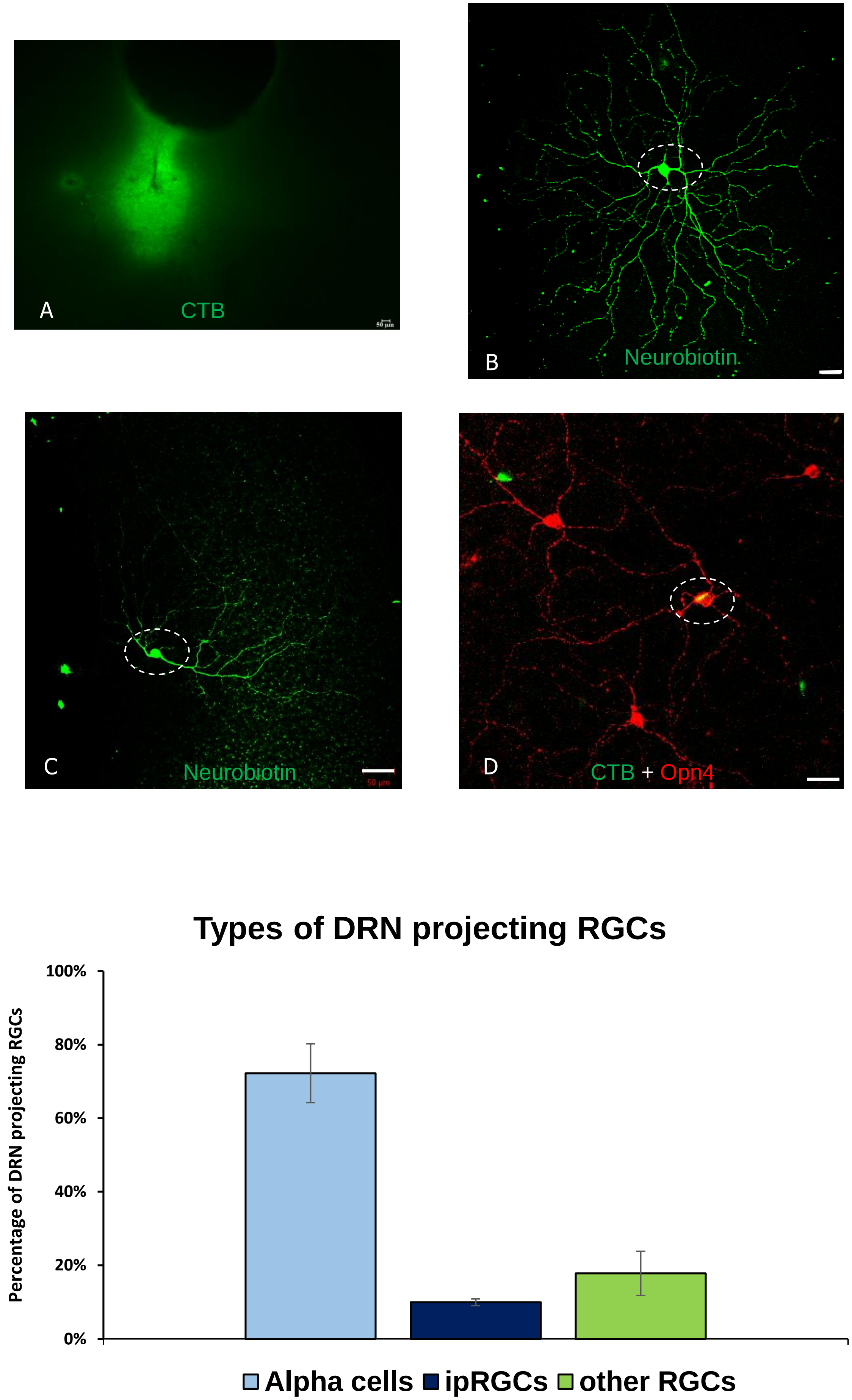
A
CTB
B
Neurobiotin
C
Neurobiotin
D
CTB + Opn4
Types of DRN projecting RGCs
Percentage of DRN projecting RGCs
100%
80%
60%
40%
20%
0%
Alpha cells
ipRGCs
other RGCs

Figure S5

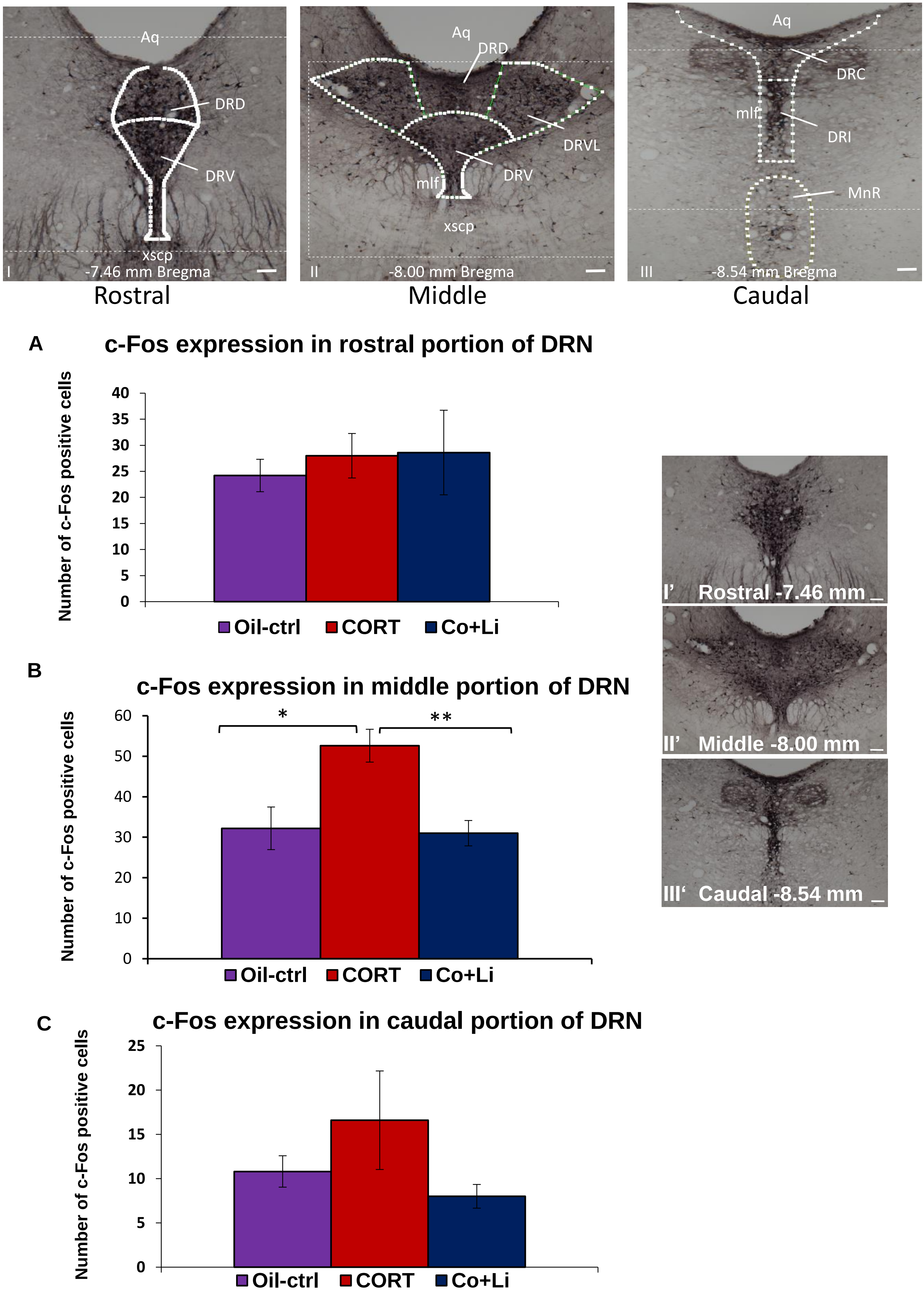

Aq
DRD
DRV
xscp
I
-7.46 mm Bregma
Aq
DRD
DRVL
DRV
mlf
xscp
II
-8.00 mm Bregma
Aq
DRC
mlf
DRI
MnR
III
-8.54 mm Bregma
Rostral
Middle
Caudal
A
c-Fos expression in rostral portion of DRN
Number of c-Fos positive cells
0
5
10
15
20
25
30
35
40
Oil-ctrl
CORT
Co+Li
B
c-Fos expression in middle portion of DRN
Number of c-Fos positive cells
0
10
20
30
40
50
60
*
**
Oil-ctrl
CORT
Co+Li
C
c-Fos expression in caudal portion of DRN
Number of c-Fos positive cells
0
5
10
15
20
25
Oil-ctrl
CORT
Co+Li
I' Rostral -7.46 mm
II' Middle -8.00 mm
III' Caudal -8.54 mm

### Figure S5

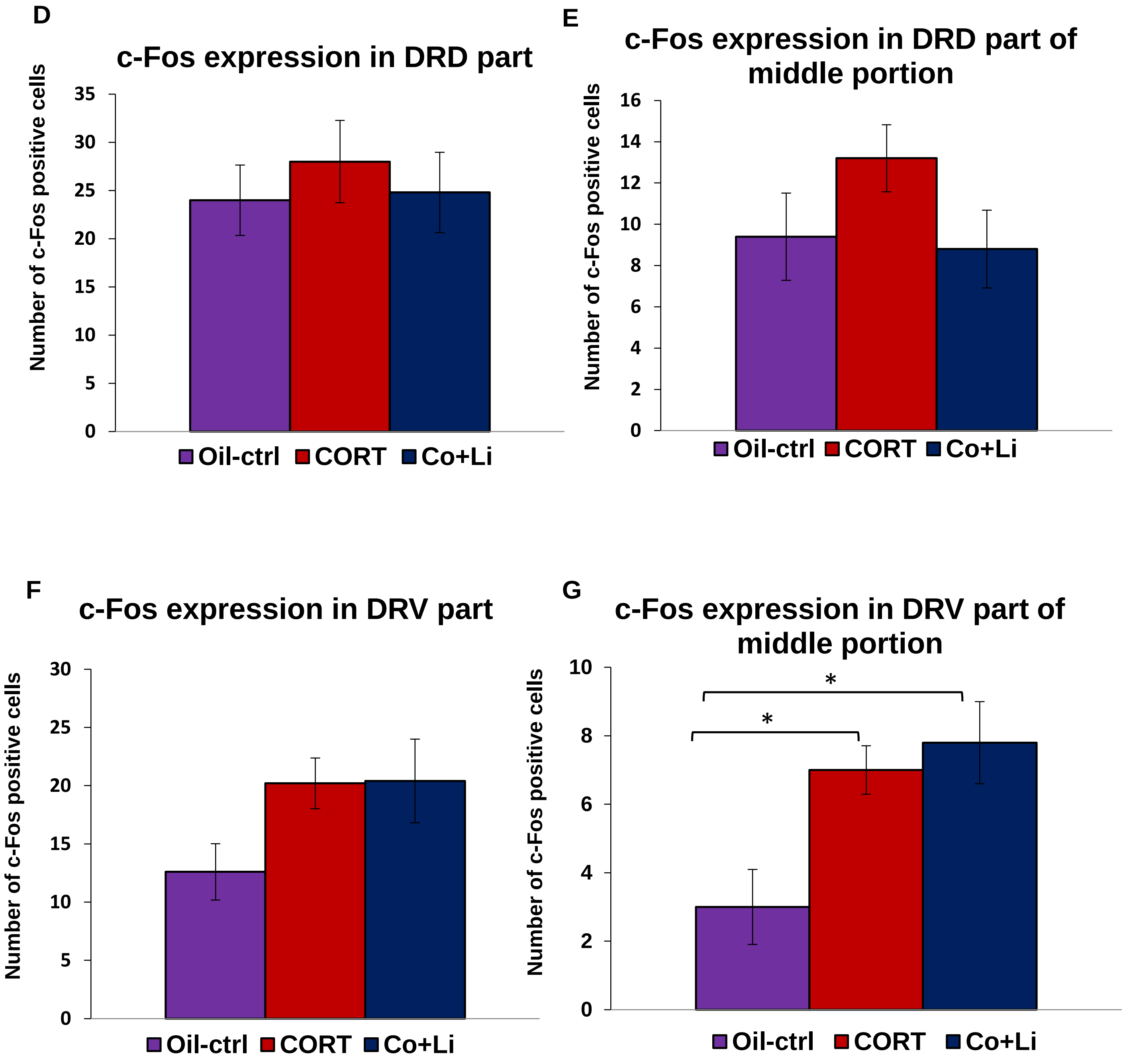
D
c-Fos expression in DRD part
Number of c-Fos positive cells
0
5
10
15
20
25
30
35
Oil-ctrl
CORT
Co+Li
E
c-Fos expression in DRD part of middle portion
Number of c-Fos positive cells
0
2
4
6
8
10
12
14
16
Oil-ctrl
CORT
Co+Li
F
c-Fos expression in DRV part
Number of c-Fos positive cells
0
5
10
15
20
25
30
Oil-ctrl
CORT
Co+Li
G
c-Fos expression in DRV part of middle portion
Number of c-Fos positive cells
0
2
4
6
8
10
*
*
Oil-ctrl
CORT
Co+Li

### Figure S6

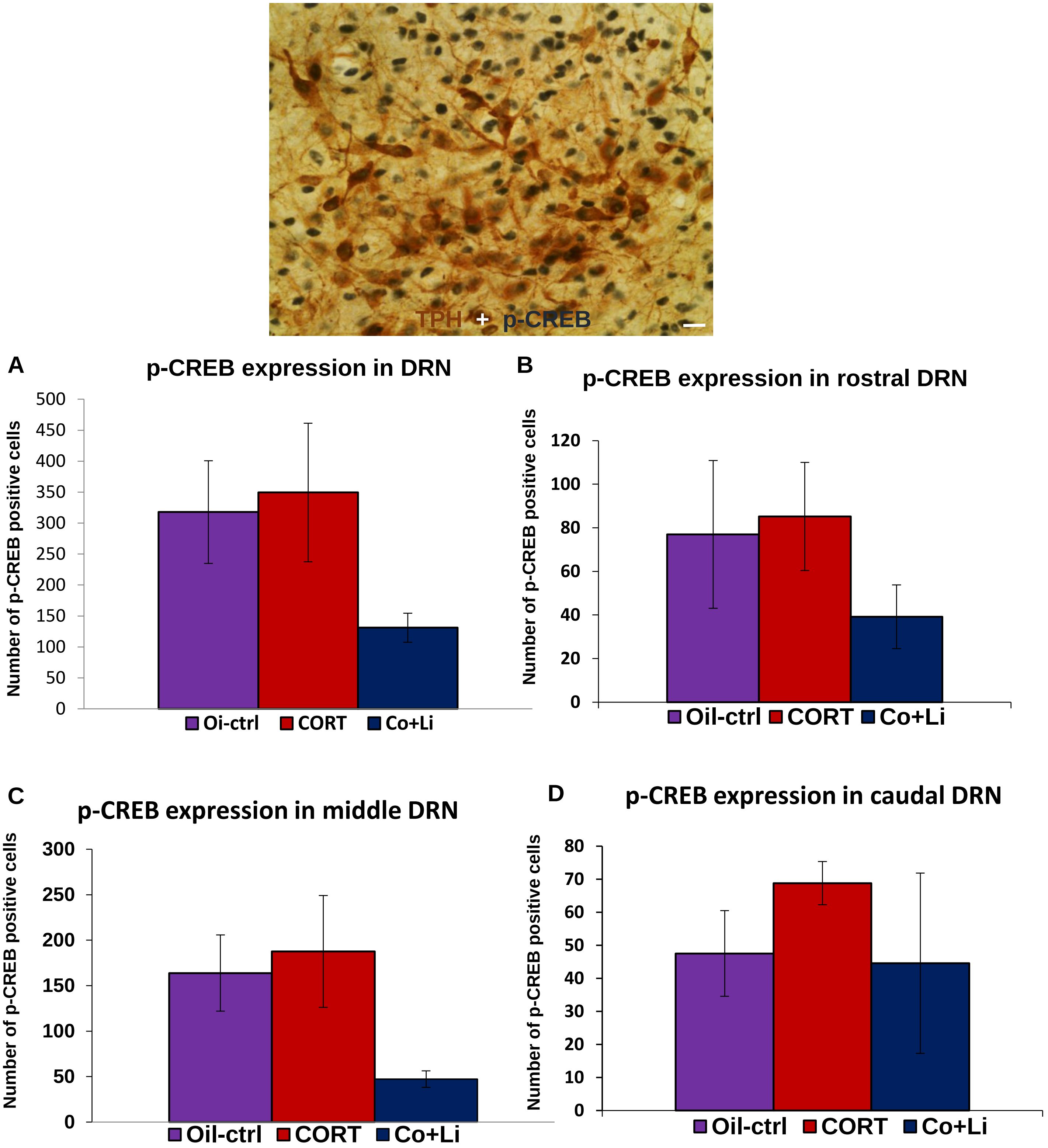
TPH + p-CREB
A
p-CREB expression in DRN
Number of p-CREB positive cells
500
450
400
350
300
250
200
150
100
50
0
Oi-ctrl
CORT
Co+Li
B
p-CREB expression in rostral DRN
Number of p-CREB positive cells
120
100
80
60
40
20
0
Oil-ctrl
CORT
Co+Li
C
p-CREB expression in middle DRN
Number of p-CREB positive cells
300
250
200
150
100
50
0
Oil-ctrl
CORT
Co+Li
D
p-CREB expression in caudal DRN
Number of p-CREB positive cells
80
70
60
50
40
30
20
10
0
Oil-ctrl
CORT
Co+Li

Figure S7

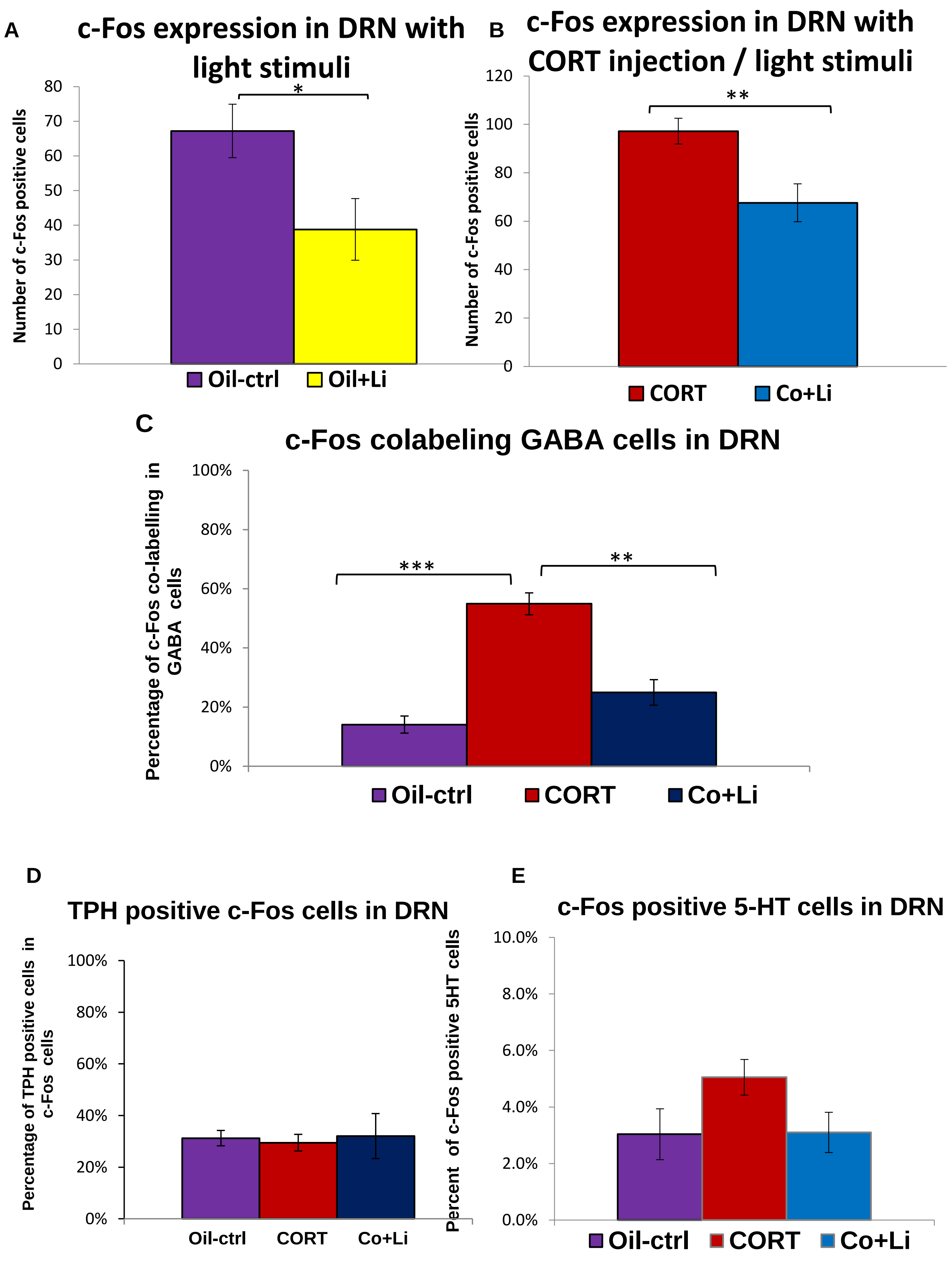
A
c-Fos expression in DRN with light stimuli
Number of c-Fos positive cells
0
10
20
30
40
50
60
70
80
*
Oil-ctrl
Oil+Li
B
c-Fos expression in DRN with CORT injection / light stimuli
Number of c-Fos positive cells
0
20
40
60
80
100
120
**
CORT
Co+Li
C
c-Fos colabeling GABA cells in DRN
Percentage of c-Fos co-labelling in GABA cells
0%
20%
40%
60%
80%
100%
***
**
Oil-ctrl
CORT
Co+Li
D
TPH positive c-Fos cells in DRN
Percentage of TPH positive cells in c-Fos cells
0%
20%
40%
60%
80%
100%
Oil-ctrl
CORT
Co+Li
E
c-Fos positive 5-HT cells in DRN
Percent of c-Fos positive 5HT cells
0.0%
2.0%
4.0%
6.0%
8.0%
10.0%
Oil-ctrl
CORT
Co+Li

### Figure S8

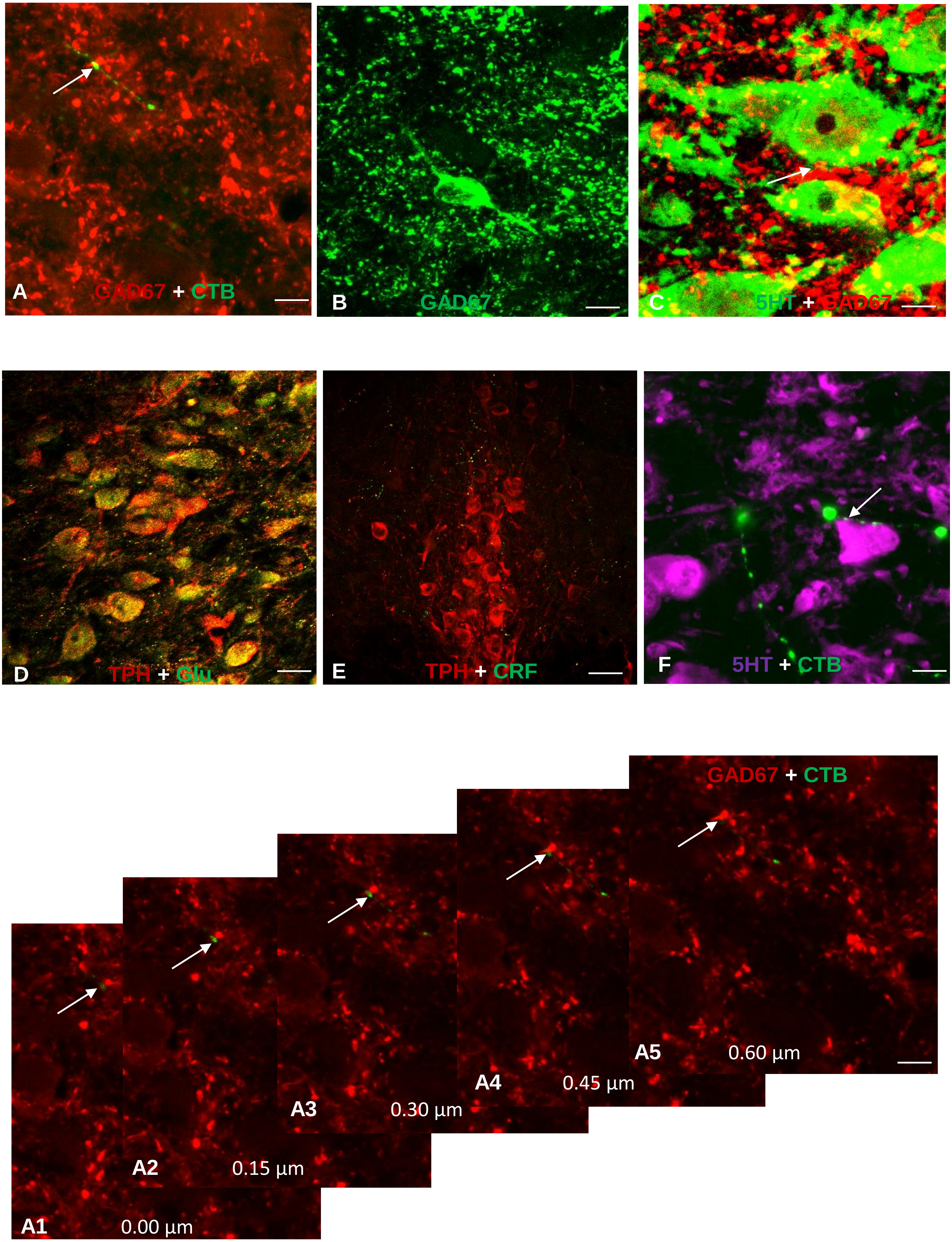
A
GAD67 + CTB
B
GAD67
C
5HT + GAD67
D
TPH + Glu
E
TPH + CRF
F
5HT + CTB
GAD67 + CTB
A1
0.00 µm
A2
0.15 µm
A3
0.30 µm
A4
0.45 µm
A5
0.60 µm

Figure S9

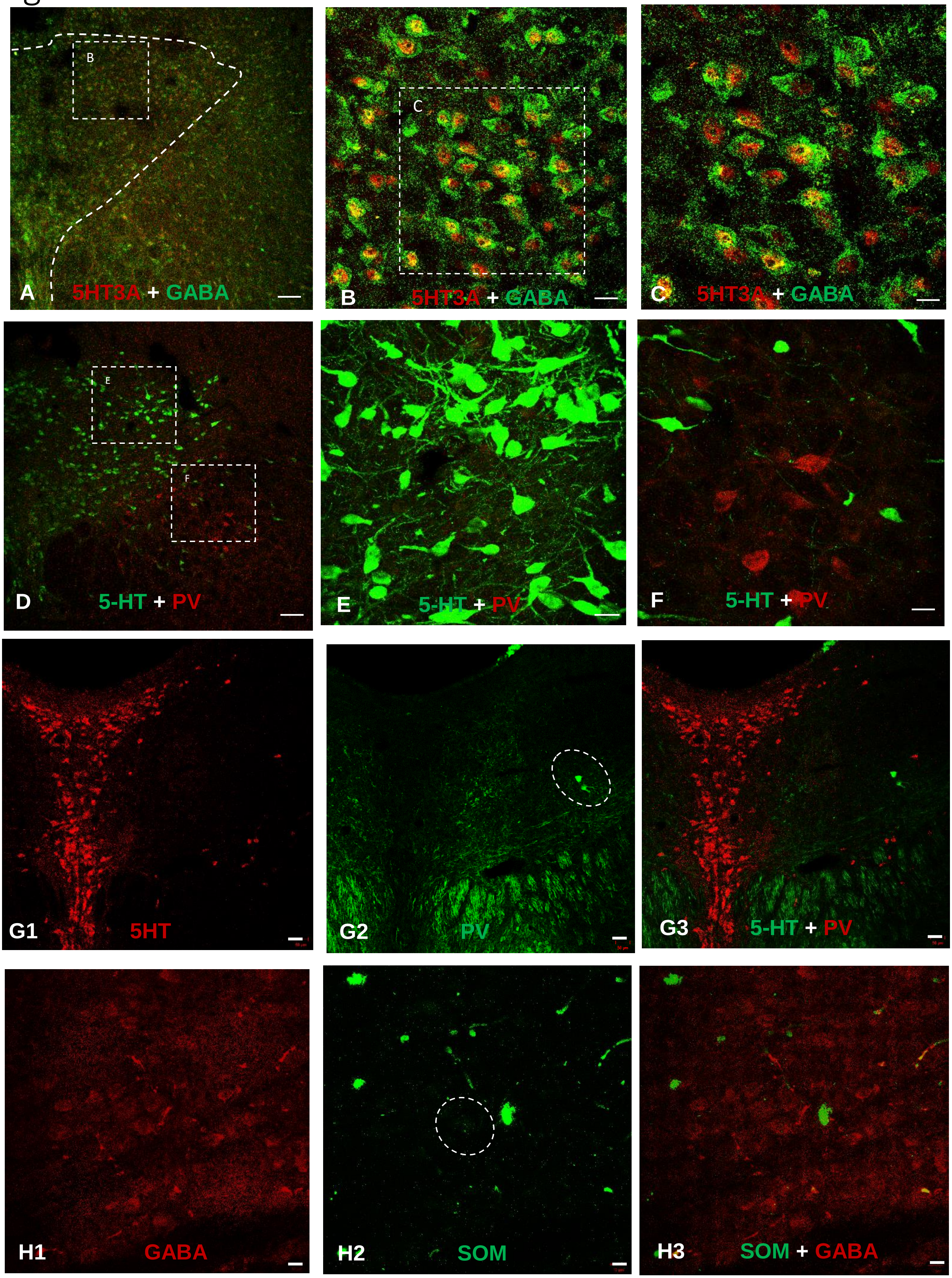
B
A
5HT3A + GABA
C
B
5HT3A + GABA
C
5HT3A + GABA
E
F
D
5-HT + PV
E
5-HT + PV
F
5-HT + PV
G1
5HT
G2
PV
G3
5-HT + PV
H1
GABA
H2
SOM
H3
SOM + GABA

**Figure S10**

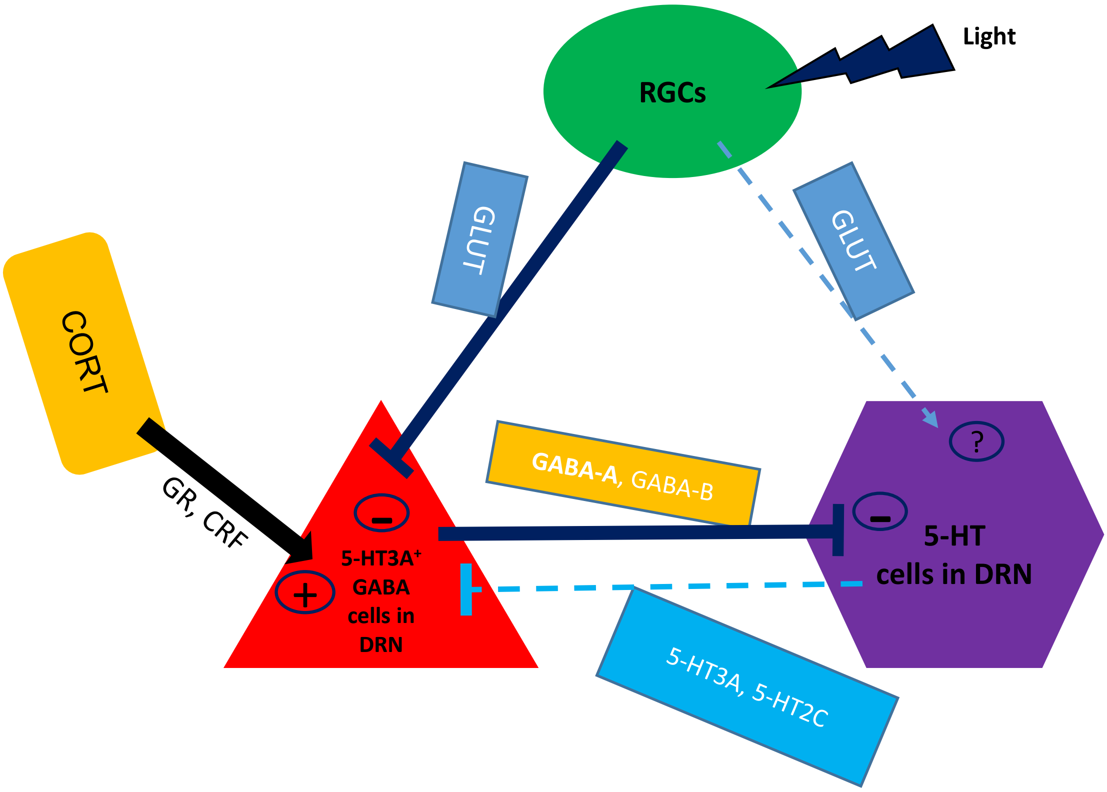
Light
RGCs
GLUT
GLUT
CORT
GR, CRF
GABA-A, GABA-B
?
5-HT3A⁺
GABA
cells in
DRN
5-HT
cells in DRN
5-HT3A, 5-HT2C
